# Insights From Protein Frustration Analysis of BRD4-Cereblon Degrader Ternary Complexes Show Separation of Strong from Weak Degraders

**DOI:** 10.1101/2025.02.09.637153

**Authors:** Tianyi Yang, Elizaveta Mukhaleva, Wenyuan Wei, Dahlia Weiss, Ning Ma, Veerabahu Shanmugasundaram, Nagarajan Vaidehi

## Abstract

PROteolysis TArgeting Chimeras (PROTACs), also known as Ligand-Directed Degraders (LDDs), are an innovative class of small molecules that leverage the ubiquitin-proteasome system to induce the degradation of target proteins. Structure based design methods are not readily applicable for designing LDDs due to the dynamic nature of the ternary complexes. This study investigates the dynamic properties of five LDD-mediated BRD4-Cereblon complexes, focusing on the challenges of evaluating linker efficiency due to the difficulty in identifying suitable computational metrics that correlate well with the cooperativity or degradation propensity of LDDs. We uncovered that protein frustration, a concept originally developed to understand protein folding, calculated for the residues in the protein-protein interface of the LDD-mediated ternary complexes recapitulate the strength of degradation of the LDDs. Our findings indicated that hydrophobic residues in the interface are among the highly frustrated residues pairs, and they are crucial in distinguishing strong degraders from weak ones. By analyzing frustration patterns, we identified key residues and interactions critical to the effectiveness of the ternary complex. These insights provide practical guidelines for designing and prioritizing more efficient degraders, paving the way for the development of next-generation LDDs with improved therapeutic potential.

## Introduction

PROteolysis TArgeting Chimeras (PROTACs), also known as Ligand-Directed Degraders (LDDs), are an innovative class of therapeutics that induce the degradation of target proteins by harnessing the ubiquitin-proteasome system^1–4^. LDD molecules facilitate the degradation of a protein of interest (POI) by recruiting an E3 ubiquitin ligase, leading to ubiquitin-mediated proteasomal degradation of the POI. These heterobifunctional molecules consist of two binding motifs connected by a linker, one binding the POI and the other binding the E3 ligase. Unlike traditional inhibitors that simply block protein function, LDDs offer a mechanism for complete target protein removal. LDDs have shown promise in treating diseases such as cancer, neurodegenerative disorders, and infectious diseases^5–7^. Recent advancements have been made in the computational modeling of the ternary complexes formed by the LDD with the POI and the E3 ligase, using molecular docking, molecular dynamics (MD) simulations, and other in silico techniques to predict the formation and stability of the ternary complexes necessary for effective protein degradation^8–10^. These studies have facilitated the identification of potential LDD candidates and have helped in optimizing their structures for enhanced efficacy and selectivity^11^.

Despite these advancements, several challenges remain in the development of LDDs. One such challenge is the impediment in using the conventional structure-based ligand design methods using crystal structures of the ternary complexes. This is because LDD ternary complexes are very dynamic and the challenge includes the design of efficient linkers that would balance the ternary complex stability with the catalytic efficiency of ubiquitin transfer. Virtual screening and structure-based drug design have played a crucial role in addressing these issues by enabling the rapid evaluation of large compound libraries and predicting the interactions between LDDs or molecular glues and their targets^11–13^. However, a fast and efficient computational tool to calculate properties that would evaluate the degradation efficiency (DC50) of LDDs ahead of synthesis is still lacking.

BRD4 (Bromodomain-containing protein 4) is a pivotal target in LDD studies due to its role in regulating gene expression by binding to acetylated histones. It is implicated in various cancers, making it an attractive therapeutic target^14, 15^. Studies have shown that LDD-mediated degradation of BRD4 leads to significant antitumor activity, highlighting its therapeutic potential^16–18^. The well-characterized nature of BRD4, along with its established interactions with E3 ligases, and the availability of several crystal structures of BRD4-LDD-Cereblon complexes, makes it an ideal system to study LDD ternary complex dynamics and its mechanisms. In this study using the BRD4-LDD-Cereblon complex structures in combination with molecular dynamics (MD) simulations we have identified computational properties of protein-protein interactions and their correlation to the degradation efficiency of LDDs.

Frustration is a concept originally developed to understand protein folding, referring to the presence of conflicting interactions within a protein that prevent it from reaching a single, well-defined energy minimum^19–21^. This concept has been extended to study protein-protein interactions (PPIs), where it helps identify regions of structural flexibility and functionality^22, 23^. In this study, using the crystal structures of the BRD4-Cereblon with 5 different LDDs as a test system^24^, we showed that residue pairs that are in a less optimized energy state or “frustrated state “in the interface between BRD4 and cereblon recapitulate the activity of the LDD in the ternary complex towards targeted degradation (DC50). By analyzing the frustration patterns within the PPI interface, we have identified key residues and interactions that contribute to the efficiency of LDD-induced degradation, that could potentially guide the design of more effective degraders.

## Results

### Structural Analysis of BRD4-Cereblon-LDD complexes

There are only five LDDs with ternary complex structures for the BRD4-Cereblon pair ^24^. In this study we have used extensive molecular dynamics simulations totaling to 6μs for each of the five LDD ternary complex to study the dynamics of the five LDD ternary complexes and calculated properties from the dynamics trajectories with the goal of being able to distinguish a strong LDD from a weak LDD. The five LDDs used in this study are shown in Fig. 1A. We refer to the five LDDs as follows: L1 (from PDB: 6BN9, dBET70, DC50 ∼ 5 nM), L2 (from PDB: 6BOY, dBET6, DC50 ∼ 10 nM), L3 (from PDB: 6BN7, dBET23, DC50 ∼ 50 nM), L4 (from PDB: 6BNB, dBET57, DC50 ∼ 500 nM), and L5 (from PDB: 6BN8, dBET55, DC50 ∼1800 nM). (https://doi.org/10.1038/s41589-018-0055-y). The degradation values are taken from literature^24^. We categorize the five LDDs into strong degraders (L1, L2, L3) and weak degraders (L4, L5) based on their DC50 values. All five LDDs have the same cereblon binder (R3 in Fig. 1A) but have different BRD4 warheads (R1 or R2 in Fig. 1A). The five LDDs differ mainly in their linker chemistry. The strong LDDs (L1, L2, L3) all have an octyl chain as the linker, but they are connected differently to the BRD4 warhead (enolate group in R1 or methyl ester group in R2) or cereblon ligand (amine or carbamate linkage). The two weak LDDs differ primarily in linker length and chemistry: L4 has a very short alkyl linker, while L5 has an eight unit ethylene glycol chain, placing them at opposite ends of the linker length spectrum. Hereafter in this paper, we will refer to the entire ternary complex as the L* complex (where * goes from LDD 1 to 5) and the LDD molecule as the L* molecule.

**Figure 1.**
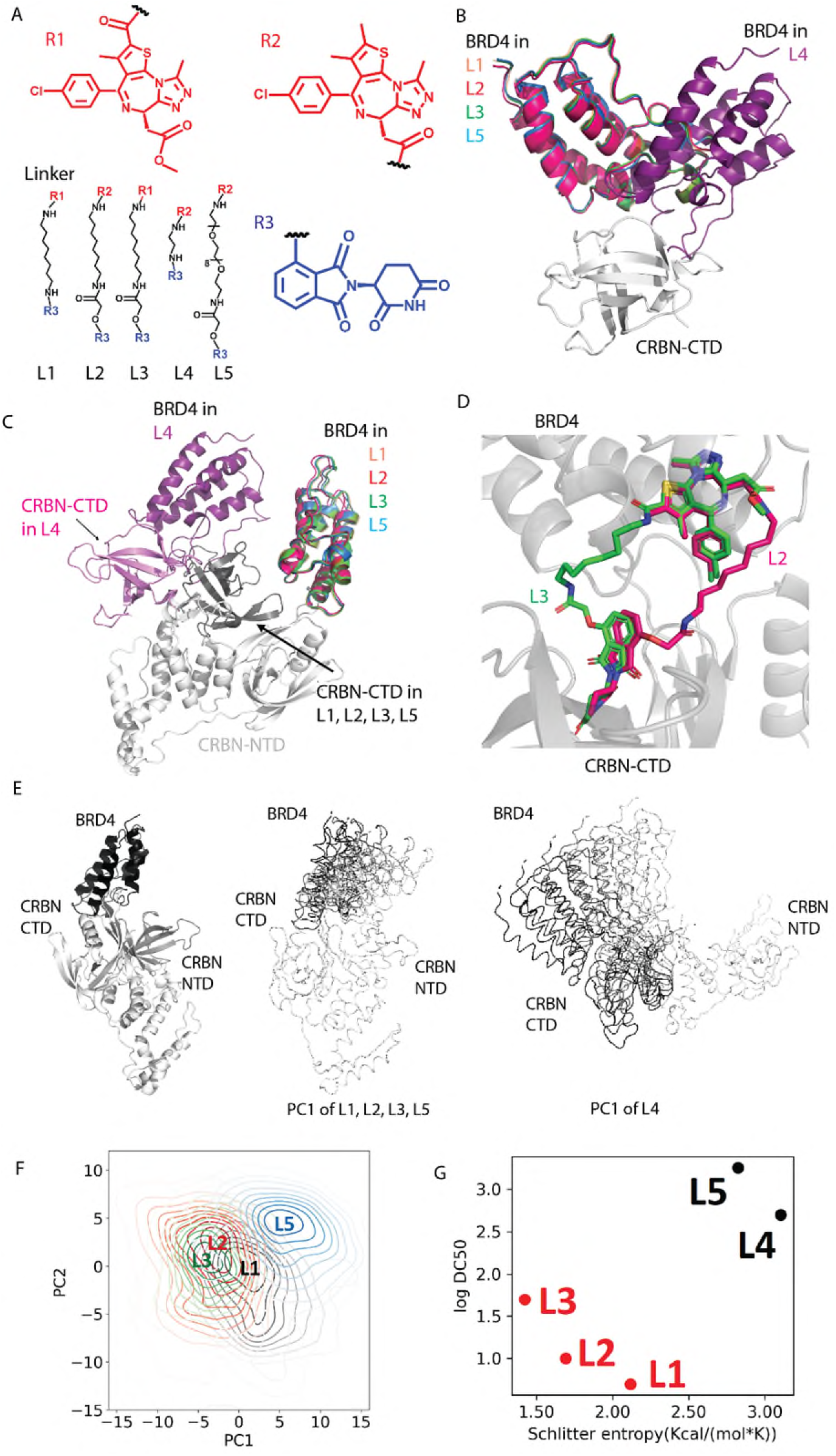
Structure and Dynamics Analysis of LDD-mediated BRD4-Cereblon Complexes. In this figure, CRBN stands for Cereblon. A) 2D structures of the five LDDs under study. B) 3D structure overlay of the five complexes, aligned by CTD. Only BRD4 and CTD are shown, with all LDDs hidden for clarity. C) 3D structure overlay of the five complexes, aligned by NTD and HBD. The whole protein is shown, with all LDDs hidden for clarity. D) 3D structure of L2 and L3 molecules in the context of the BRD4-CTD interface. E) Five equally spaced conformations in the PC1 space extracted from principal component analysis (see Methods) of the aggregated trajectories L1, L2, L3, and L5 (middle panel) and L4 (right panel). The perspective of the protein structure is shown in the left panel. F) The contour map of the conformational ensemble of all the LDD complexes projected in the PC1 and PC2. G) Plot of Conformational entropy versus logDC50 for the five LDD complexes.

Because of the different linker chemistry, there are observable differences in the LDD ternary complex structures. In this manuscript, we only considered the residues that are resolved in all the complexes of the BRD4 domain, cereblon C-Terminal Domain (CTD), N-Terminal Domain (NTD), and Helical Bundle Domain (HBD)^24^, ensuring that all systems have the same number of amino acids and comparable. Aligning the crystal structures of all the complexes by the CTD of cereblon, we observed a distinct orientation of BRD4 in L4 complex that is different from the rest of the complexes (Fig. S1). While L1, L2, L3, and L5 complexes have a highly similar CTD-BRD4 orientation, in L4 complex, the BRD4 flips to a different direction (Fig. 1B and Fig. S1). When aligned by the HBD and NTD of cereblon, we notice that the CTD in L4 complex is positioned very differently compared to the other four LDDs. The CTD in L4 complex is far away from the NTD (Fig. 1C). This could be due to the shorter linker in L4.

It should be noted that only the L2 and L3 complexes have the LDD molecule resolved in the protein-LDD complex structures (Fig. 1D). In contrast, L1, L4, and L5 complex only have the proteins resolved, but the LDD molecules were not resolved in the X-ray structure. To prepare the structures for the dynamics study, we modeled the unresolved LDD molecules into the resolved protein complex. We modeled just the LDDs, L4 and L5 molecules using L2 molecule as a template, and L1 molecule using L3 molecule as a template (see Methods), because they share the same BRD4 warhead (Fig. S2). Additionally, the modeled systems and the template systems show only minor differences in the root mean square deviation (RMSD) in coordinates in the warhead binding region (Fig. S3). This indicates that even if the LDD molecules are not resolved in the modeled system, the warheads should bind in a similar manner; otherwise, we would expect a highly different warhead binding region.

### Dynamics and Flexibility of LDD-Mediated BRD4-Cereblon Complexes

Due to the different conformation of the L4 complex, many computational metrics can easily distinguish L4 from L1, L2, and L3. However, the L5 complex is highly similar to the strong LDD complexes, making it particularly challenging to distinguish L5 from L1, L2, and L3. However, the L5 complex is like the strong LDD complexes, making it a major challenge to separate L5 from L1, L2, and L3. Starting from the crystal structures of the ternary complexes with L1 to L5 we performed 5 runs of MD simulations totaling to 6 μs for each system. To understand the domain motion in the dynamics of these complexes we performed Principal Component Analysis (PCA) (see Methods for details). The dynamics of LDDs and LDD-mediated BRD4-cereblon ternary complexes exhibit distinctly different motion in the MD simulations. Overall, the L4 complex shows a very distinct global motion compared to the other four complexes. The dominant motion in the L4 complex is the relative motion between BRD4-CTD and NTD-HBD domains (Fig. 1E right). In contrast, in the other four LDD complexes, the dominant motion occurs between BRD4 and cereblon (Fig. 1E middle). L4 makes contacts only with the CTD and not the NTD domain of cereblon and hence does not stabilize the closed domain conformation of cereblon. Therefore, the flexibility of the L4 bound ternary complex primarily arises from the relative domain motion between the NTD-CTD domains within cereblon.

The flexibility of the L4 complex is higher than that of the other four LDD complexes, as evidenced by higher RMSD in coordinates (Fig. S4) and a wider spread of the conformations in the PC space (Fig. S5). The L5 complex also shows distinctly displaced ensemble of conformations compared to L1, L2 and L3 complexes in the PC space perhaps due to the long flexible linker (Fig. 1F). Focusing on the LDD molecules themselves, the L4 molecule has the lowest flexibility as seen in the RMSD versus time plots, due to its short linker, while the L5 molecule has the highest flexibility due to its longest linker (Fig. S6). We then calculated the conformational entropy using the populations of the different microstates in the PC landscape. The conformational entropy calculated from the populations of the various conformations (see Methods), cannot distinguish strong and weak LDD complexes (Fig. S7). However, the entropy of the residues in the BRD4-CTD protein-protein interface can clearly separate strong from weak LDD complexes (Fig. 1G). This indicates that protein interface plays a critical role in recapitulating the strength of LDDs rather than the overall properties of the ternary complexes.

### Evaluating Protein Frustration in LDD-Mediated BRD4-Cereblon complexes

Since the properties of the protein-protein interface (PPI) in the ternary complexes more closely recapitulate the effectiveness of the LDDs we turned our attention to studying the interface more in detail. Protein frustration is an increasingly recognized concept when discussing PPIs^25^. Originally developed to study protein folding, frustration occurs when residue interactions within a protein or between proteins are in a sub-optimal energy state. Previous studies have shown that although the overall protein might be in an energy optimal state, structural regions and/or residue pairs might be in a sub-optimal energy state. Such “frustrated residue pairs” in a protein structure are typically in the active site or on the surface of proteins where other proteins couple and form complexes ^20^. Recent work has shown that residues in the ligand binding sites can be frustrated^23^. In this study, to understand how frustration relates to effectiveness of LDDs, we calculated the mutational frustration^26^ for every residue pair using the Frustratometer^27^ software for the crystal structures of the ternary complexes and also for all the MD snapshots from the trajectories of the 5 ternary complexes.

Frustratometer classifies residue pairs as highly, neutrally, or minimally frustrated based on pair-wise interaction energies compared to mutations at the same position. For a residue pair, if the wild type (WT) energy is more favorable than most mutations (with a Δ(WT-Mut) Z-score less than −1), the pair is considered minimally frustrated. Conversely, if the WT energy is less favorable than most mutations (with a Δ(WT-Mut) Z-score greater than 1), the pair is highly frustrated. If the WT energy is close to the average of all mutations (with a Δ(WT-Mut) Z-score between −1 and 1), the pair is neutrally frustrated.

We calculated the number of highly frustrated residue pairs in the POI-E3 interface in the ternary complexes. For each LDD complex, we calculated the percentage of the highly frustrated residue pairs to the total number of residue pairs in the BRD4-cereblon interface (Fig. 2A), for each MD simulation snapshot. The interface residue pairs come from both inter-protein (BRD4-cereblon) or intra protein (within BRD4 or within cereblon, but both residues of the pair are in the interface). We then calculated the average of this percentage over MD snapshots. This is defined as the frustration level.

**Figure 2.**
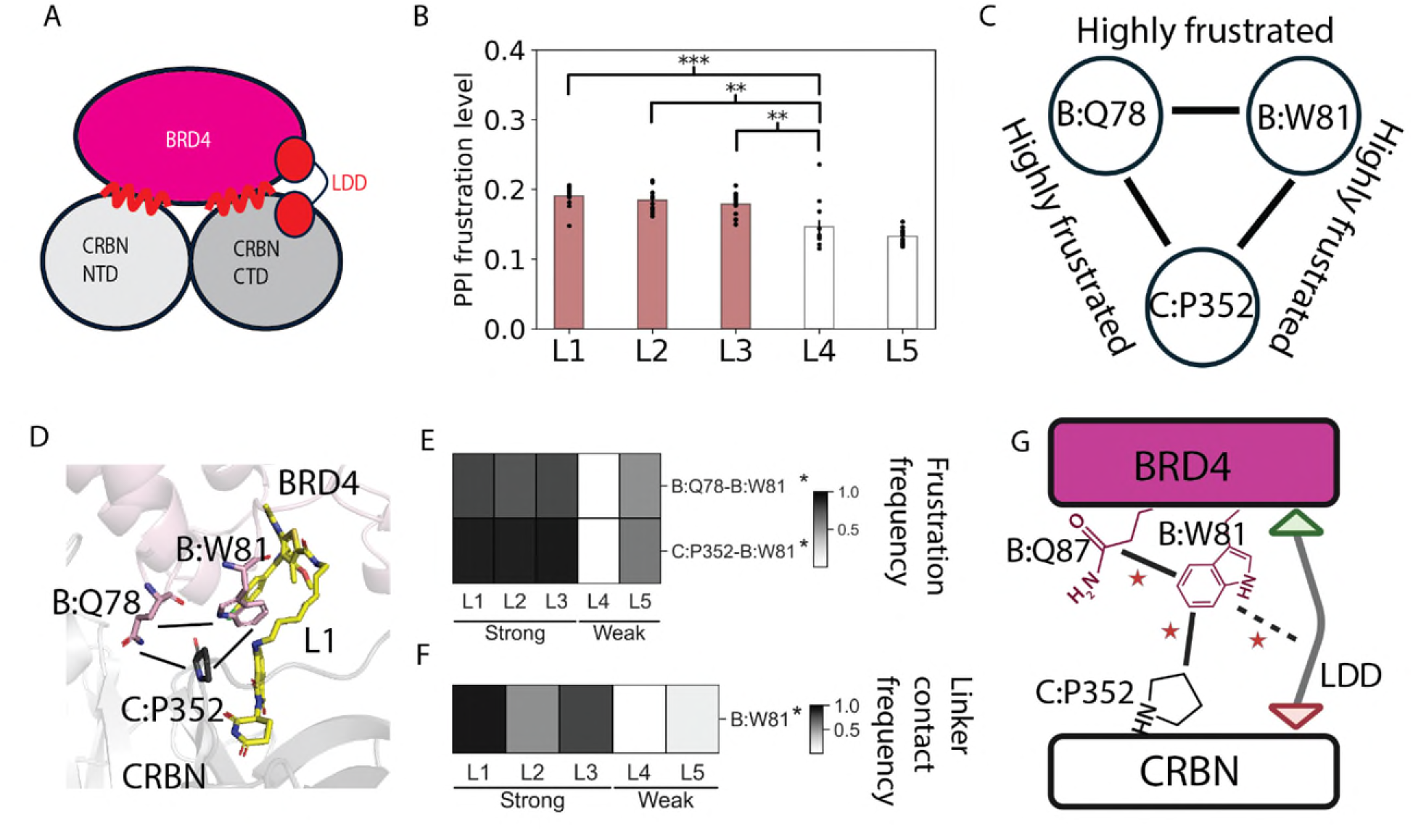
Frustration in the BRD4-LDD-Cereblon complex. In this figure, CRBN stands for Cereblon. A) A 2D scheme to show the PPI region (red curve) considered for frustration calculation in this study. B) PPI frustration level comparison among the five LDDs complex. P-value for significance: L1-L4: 0.0003; L2-L4: 0.001; L3-L4: 0.004. L1-L5: 3.8e-10; L2-L5: 4.3e-9; L3-L5: 6.9e-8. C) The top 3 persistent (in terms of frequency in MD simulations) highly frustrated contact pairs in L1 shown as Cartoon. D) The top 3 persistent (in terms of frequency in MD simulations) highly frustrated contact pairs in L1 mapped on the 3D structure. The short black lines show the persistent highly frustrated contact pairs. BRD4’s cartoon and sidechains are in pink. Cereblon’s cartoon and sidechains are in grey. E) Heatmap of frequencies of highly frustrated residue pairs in the BRD4-cereblon interface and between PPI residue pairs. Pairs that are shown here are those with significant difference (p value<0.05) between strong and weak LDD system and frequency greater than 70% in at least one system. F) Contact frequency between LDDs’ linker and protein residues, with contact frequency greater than 70% in at least one system. B stands for BRD4. G) Proposed mechanism scheme to explain how LDD linker affects PPI frustration. Dashed line refers to LDD linker-protein residue contact. Solid lines refer to protein residue frustration.

As shown in Fig. 2B, the PPI frustration level clearly separates strong LDD complexes from weak ones. The strong LDDs (L1, L2, and L3) show a more frustrated PPI compared to weak LDDs (L4 and L5). This suggests that frustration values at the PPI calculated using a conformational ensemble is a computational metric that can be used to sort strong LDDs from weak LDDs prior to synthesis, in cases with a high confidence model of the ternary complex.

For an unknown set of degraders, we came up with a thumb rule to distinguish the strong ones from the weak. We plotted the distribution of the number of trajectories that are within a certain frustration level range in the PPI as shown in Fig. S8. The gaussian kernel density fit shown in black in the figure, contains three peaks. The boundary between weak and strong is ∼0.4 for this BRD4:CRBN system. However, this cutoff value in frustration level may not apply for other target protein or E3 ligase ternary complexes. For an unknown case, we propose that the LDDs under the tallest peak on the high frustration level are predicted to be strong and can be prioritized for testing.

To identify the location of highly frustrated interactions within the system, we mapped the top three persistently highly frustrated residue contacts (based on the number of MD snapshots that they showed frustration in, across multiple MD simulations) onto the structures (Fig. 2C, D, Fig. S8). The 3 strong LDD systems (L1, L2, L3) share the same top 3 persistently highly frustrated pairs, which are C:P352-B:W81, C:P352-B:Q78 and B:Q78-B:W81. In contrast, the two weaker LDD systems show variations. In L4 ternary complex, BRD4 binds to cereblon in a different orientation compared to the other four LDD systems, resulting in a different set of top three persistently highly frustrated contacts: B:K91-B:N93, C:S396-B:N93 and C:Q390-C:S396. The L5 ternary complex, however, shows a binding conformation like the three strong LDD systems and shares two of the top three persistent highly frustrated pairs with them: P352-W81 and P352-Q78. The third pair for L5 is B:K155-C:G151. To quantitatively assess the protein-protein interaction residues that distinguish between strong and weak LDD complexes, we calculated the p-value of the frequency of highly frustrated residue interactions between the strong group (L1, L2, L3) and the weak group (L4, L5) for each contact pair. Two persistent interactions (frequency > 70% in at least one system) were identified as significantly different, with a p-value less than 0.05, as shown in Fig. 2E. One residue pair, C:P352 and B:W81 (C indicates from cereblon and B for BRD4), is highly frustrated in strong LDD complexes (82%∼86%) and less so in weak LDD complexes (60% in L5 complex). It should be noted that although C:P352 and B:W81 is in the top 3 persistently frustrated residue pairs in L5, its frequency is still far less (61%) in the L5 LDD complex compared to strong LDD L1 (89%), L2 (89%), L3 (91%) complexes. Another pair, B:W81 and B:Q78, may also potentially distinguish the two categories but with less separation (85%∼90% in strong LDD complexes and 74% in L5 complex). In the 3D structure (Fig. 2D, Fig. S8), we observed that B:W81 is sandwiched between the cereblon ligand and the BRD4 warhead of the LDDs, while C:P352 and B:Q78 are immediately around this region.

To further explore the impact of LDD molecules on the protein interface, we calculated the contact frequency—defined as the percentage of MD simulation snapshots where two residues are in contact—between the LDD and the protein. The contacts involving the two binding moieties of the LDD (Fig. S10A, B) and the linker of the LDD (Fig. 2F) were analyzed separately. On the contact heatmaps, only residues showing significant differences in contact frequency between strong and weak LDD systems are displayed. For residues that are in contact with LDD’s binding moiety (Fig. S10A, B), B:F83, B:V87 and C:N351 exhibit more persistent interactions with strong LDDs compared to weak ones. The heatmap for LDD linker-protein contacts (Fig. 2F) indicates that W81 in BRD4 consistently contacts the linker in strong LDD complexes, but to a lesser extent in weak LDD complexes. Combination of frustration analysis and LDD-protein contacts, shows that the linker of strong LDDs form more stable interactions with W81 on BRD4 compared to weak LDDs. This suggests that strong LDDs may lead to more persistently highly frustrated pairs of C:P352-B:W81 and B:Q78-B:W81 than observed in weak LDDs(Fig. 2G).

### Differentiating Strong and Weak LDD Complexes Through Hydrophobic Patch Integrity and Water Energy Profiles

The residue contact frequency in the PPI (Fig. S10C), shows significant difference between strong and weak LDDs. There are several hydrophobic residues located in the cereblon:BRD4 interface, including B:L148, B:M149, C:F102, C:F150, B:F79, C:P352 (Fig. S10C). These hydrophobic residues form a tight hydrophobic patch in the L1 ternary complex as shown in Fig. S10D. The results show the critical role of hydrophobic residues and their interactions in distinguishing strong from weak LDDs. This suggests that water molecules may significantly influence PPIs, potentially affecting the efficiency of LDDs as detailed in this section.

To evaluate the effect of the entry of water into the PPI and LDD binding site, we calculated the water occupancy map using the VolMap tool from VMD (see Methods). As shown in Fig. 3, the hydrophobic patch of residues formed by F102 and F150 from cereblon with F79 from BRD4 shows no persistent water molecules during MD simulations. Additionally, there is another hydrophobic patch around residue W81 in BRD4 with the aromatic groups of the cereblon ligand and the BRD4 warhead (Fig. 3A left). The same trend is observed in the L2 and L3 complexes (Fig. S11A). However, in the L5 complex, although a similar hydrophobic patch F102(cereblon) - F79 (BRD4)-F150 (cereblon) exists, the hydrophobic patch formed by the W81 with the aromatic moieties of the LDD is broken (Fig. 3A right). In the L4 complex, there is no hydrophobic patch observed around the LDD linkers, and no hydrophobic patch formed by F102 (cereblon)-F79(BRD4)-F150(cereblon) due to the different orientation of BRD4 (Fig. S11A). For L4, the previously described difference in the orientation of BRD4 with respect to cereblon prevents the formation of the same hydrophobic path observed in other complexes. Instead, the L4 complex stability relied on the short linker and surrounding residues. In short, the weak L4 and L5 LDD complexes have less hydrophobic residues in PPI, while the strong LDD complexes have two hydrophobic patches in PPI.

**Figure 3.**
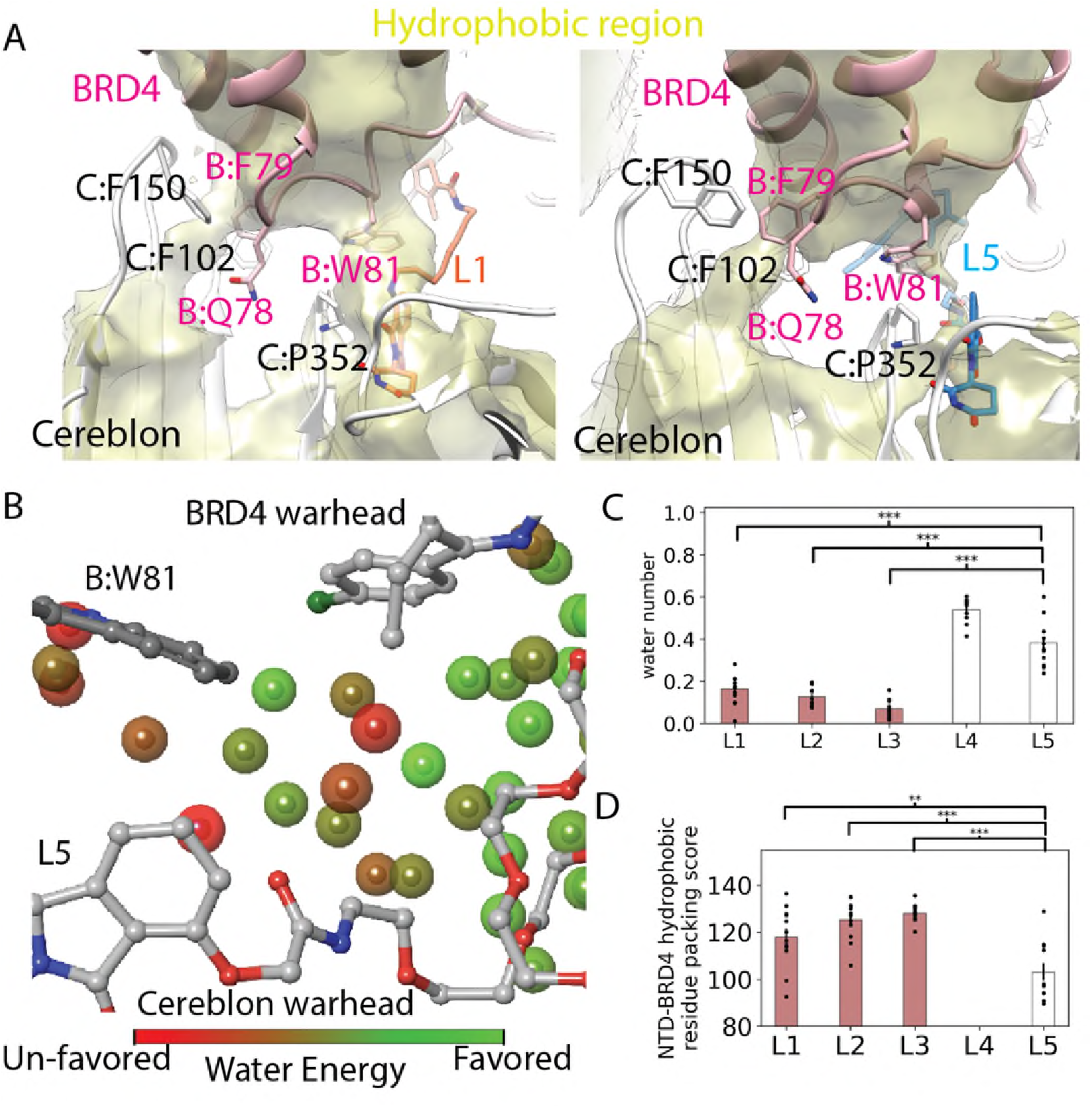
Water plays a critical role in the interface of the ternary complexes. A) Water occupancy <1% over simulations in the Hydrophobic region for L1 (left) and L5 (right). B) Water energy for one frame from the L5 simulation, zooming in on the highlighted water energy around W81. C) Counting the number of waters around W81, using 3 Angstrom as the cutoff. P-value for significance: L1-L4: 1.1e-10; L2-L4: 2.3e-15; L3-L4:5.4e-17. L1-L5: 2.1e-5; L2-L5: 1.9e-7; L3-L5:2.7e-10. D) Residue-Residue packing score (RRCS) of the hydrophobic core formed by C:F102-B:F79-C:F150 hydrophobic patch. P-value for significance: L1-L5: 0.004; L2-L5: 2.8e-5; L3-L5: 8e-7.

Is water within these hydrophobic patches energetically unfavorable? To answer this question, we calculated the water energy using the Maestro water map module (see Methods for details). The results confirm that water molecules around hydrophobic patches have unfavorable energy (shown as red spheres Fig. 3B). We counted the total number of waters in both hydrophobic patches and found significantly more water around B:W81 in the L4 and L5 complexes (Fig. 3C), which could distinguish weak LDD complexes from the strong ones. However, the number of water molecules in the F102-F79-F150 hydrophobic patch cannot significantly separate the L5 complex from strong LDD complexes (Fig. S11B), indicating that even in the weak LDD complex, water will not penetrate the hydrophobic core and destabilize the PPI.

Although there is no significant difference in the water numbers in the F102-F79-F150 hydrophobic core, the residue packing in this hydrophobic core is weaker in the L5 complex (Fig. 3D). We analyzed the packing between residues in BRD4 and cereblon using a program called Residue-Residue Contact Score (RRCS)^28^. RRCS evaluates the packing efficiency between two residues by summing up the distances between every possible pair of non-hydrogen atoms from the two residues. This difference in packing suggests that the F102-F79-F150 hydrophobic patch is indeed less well-packed in the L5 complex, although not to the degree that would allow water penetration. Based on the analysis of the water profile, we conclude that strong and weak LDD complexes can be separated based on their ability to maintain the hydrophobic patches in the PPI.

## Discussion

Our data suggest a confluence of multiple mechanisms underlying the degradation efficiency of LDDs. A strong LDD with high degradation efficiency must maintain a proper hydrophobic interface between BRD4 and cereblon, while a weak LDD with lower degradation efficiency may exhibit a less hydrophobic PPI due to various factors as illustrated in the scheme in Fig. 4.

**Figure 4.**
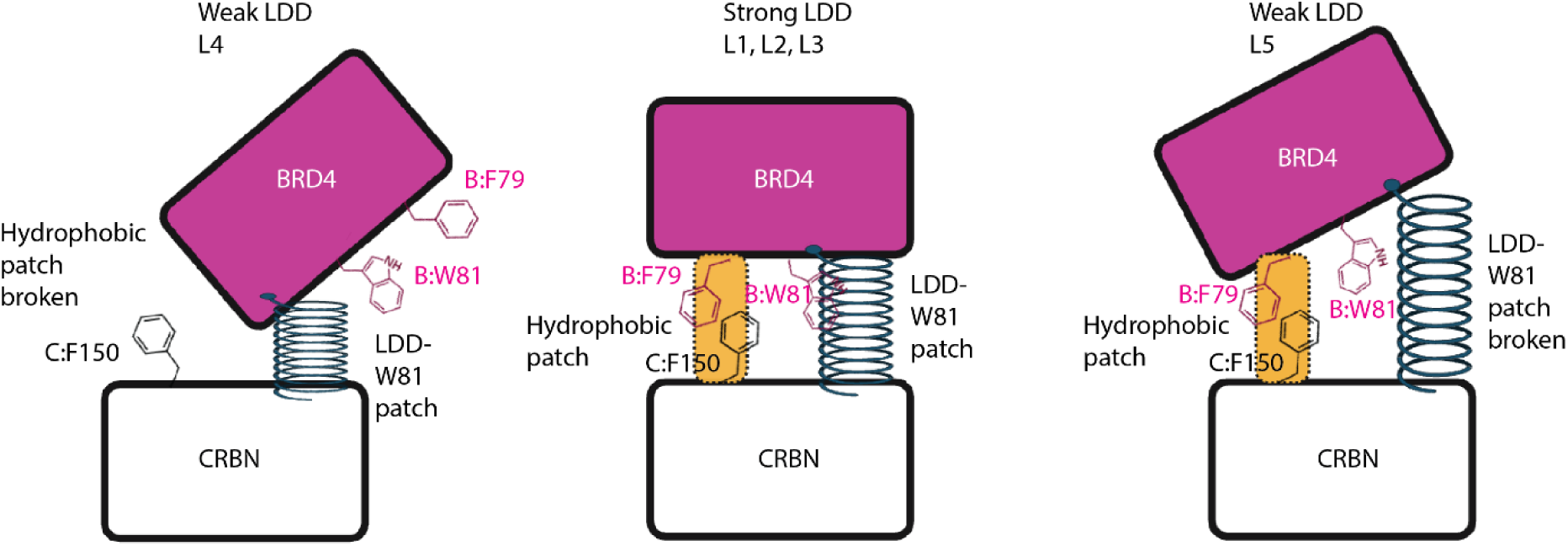
Proposed mechanism scheme to explain the degradation efficiency of LDDs. CRBN stands for Cereblon.

The linker allows for proper rotation and flexibility, enabling the warheads and E3 ligase ligand to pack with the ring of W81 forming the ‘LDD-B:W81 hydrophobic patch’. Additionally, the linker must be sufficiently extended so that BRD4 and cereblon can form the ‘C:F102-B:F79-C:F150 hydrophobic patch’. If the linker is not extended enough and lacks flexibility, as seen in the L4 complex, it forces BRD4 into an orientation relative to cereblon that causes C:F102, C:150, and B:F79 to point in different directions, thus breaking the C:F102-B:F79-C:F150 hydrophobic patch. Consequently, the BRD4-cereblon PPI is primarily maintained by the LDD linker and a few surrounding residues, leading to a worse DC50.

Conversely, if the linker is too extended and very flexible, as in the L5 complex, while the C:F102-B:F79-C:F150 hydrophobic patch can still form, albeit with less stability, the linker is too flexible to pack B:81 between the two warheads, thus disrupting the LDD-B:W81 hydrophobic patch. As a result, the BRD4-cereblon PPI is mainly maintained by the C:F102-B:F79-C:F150 hydrophobic patch and lacks support from the LDD-B:W81 hydrophobic patch, also leading to a worse DC50.

In summary, our work has identified that residue pair frustration in the interface cereblon and the protein of interest recapitulates the degradation efficacy of LDDs. This metric is new to the LDD field and can be used to prioritize synthesis of multiple LDDs. By highlighting the importance of hydrophobic patches and their role in maintaining proper cereblon-BRD4 interactions, we provide valuable insights into the structural and dynamic requirements for effective LDDs. This study not only advances our understanding of LDD-mediated degradation but also offers practical guidelines for designing more efficient degraders in future research. These findings pave the way for developing next-generation LDDs with improved therapeutic potential.

## Methods

### Structure preparation and simulation setup

The cereblon-LDD-BRD4 ternary complexes crystal structures were downloaded from the protein data bank (PDB) (ID: L1 - 6BN9, L2 - 6BOY, L3 - 6BN7, L4 - 6BNB, L5 - 6BN8). To save computational time, DDB1 chain was deleted. cereblon’s residue 44-63 and 424 to 427 were deleted. Because in 6BNB, these segments are not resolved. The missing residue on cereblon (210-218) were modeled using 8CVP as template. For 6BN9, 6BNB and 6BN8, the LDDs are missing in the crystal structure. To model the ligand, 6BN7 was used as template for 6BN9. 6BOY was used as template for 6BNB and 6BN8. The template was selected based on the chemical similarity of LDD. The ligands were modeled by overlaying protein and rebuilding the linker of the LDD. Minimization of the ligand was performed in Maestro after modeling. The ternary complex was prepared using the Maestro Protein Preparation Wizard module. Capping groups were added on the termini. SelenoMET was converted to MET. For 6BN8, BRD4-A145 was mutated back D145. (it is D145 in other 4 structures).

LDD molecules were saved in mol2 format for parameterization. Minimization of the LDD was performed using MacroModel in Maestro using OPLS4 as forcefield. The PRCG^29^ method and 0.05 convergence threshold were used for minimization. LDD forcefield parameters were derived using the Antechamber module from AmberTools23^30^. Partial charges (Atomic electrostatic potential charges) were calculated using the Jaguar^31^ module in Maestro. For Jaguar, HF and 6-31G** basis set were used. Accuracy was set to ultrafine. ESP was selected for output. PBF was used for solvation model and water was selected for solvent. The charges generated in the last step of the out file were used to replace the charge in .lib file generated by Antechamber.

FF14SB^32^ force field was employed for protein. Gaff2^33^ was loaded for ligand. ZAFF^34^ forcefield was used for the coordinating Zn ion in cereblon. The ternary complex was solvated in a truncated octahedron TIP3P water box with at least 15 Å margin from the protein surface using tleap from the AmberTools 23. Na+/Cl- were used as counterion s for neutralization. Na+/Cl- were also added to make water box salt concentration of 0.15M. In Amber the N-termini cap is named NME. Chlorine is named Cl. They should be modified manually if it is not named in this way.

The MD simulations were run with Amber22^35^. A two-phase minimization procedure was performed initially. During the first phase, a minimization of 2000 steps was performed, where a constraint of 10 kcal/(mol* Å^2^) was imposed on all atoms except for water. In the subsequent phase, all atoms were allowed mobility for another 2000 steps minimization. For each round of minimization, the steepest descent method was employed for the first 1500 steps, then conjugate gradient method was used for 500 steps. The equilibration phase began with a short NVT heating stage of 0.5 ns, gradually increasing the temperature from 0K to 300K, while maintaining a 10 kcal/(mol* Å^2^) restraint on the protein and LDD. The system then underwent an eight-steps NPT equilibration process at 300K and 1 bar, during which restraints were progressively relaxed from 10 kcal/(mol* Å^2^) down to 1 kcal/mol in decrements, with each step lasting 5 ns. After this, a 50ns equilibration of the ligand occurred, by releasing the restraint on LDD but maintaining a 1 kcal/(mol* Å^2^) restraint. Then the whole system was equilibrated for 10 ns with no restraint. The equilibration’s last frame served as the starting point for the production simulations. 20 independent production runs were initiated with random velocities, extending for 500 ns each, with snapshots saved every 100 ps. Trajectories were also saved every 500ps for the following analysis. Temperature and pressure throughout the simulations were regulated using the Berendsen method^36^. The reference temperature and pressure of production run was the same as NPT equilibration. The particle Mesh Ewald method^37^ calculated the van der Waals forces. Non-bound interactions were assessed using a cutoff of 10 Å. SHAKE^38^ was applied on bonds involving hydrogen.

### Simulation analysis

#### RMSD and trajectory selection

CTD-BRD4 Cα RMSD was calculated to select trajectories that maintained stable ternary complex. For each LDD, 12 Trajectories whose RMSD running average was always lower than 4 Å were selected for the following analysis. 12 is the minimum number of trajectories that satisfied the RMSD cutoff among all the 5 LDD systems.

### Principal component analysis and conformational entropy

Principle component analysis (PCA) was performed to investigate conformational dynamics of the protein complex. The simulation trajectories were first converted to Gromacs format. Protein backbone PCA was conducted with Gromacs2022^39^. The contour map of PC1 and PC2 was plotted by concatenating trajectories of all the LDDs. Conformational flexibility was evaluated by computing Schlitter entropy^12^. For each LDD independent runs were concatenated for calculation. The Schlitter entropy was computed with Gromacs2022^40^. For entropy of the protein complex all the Cα atoms were selected for the calculation. For interface entropy, all the atoms of the residues which have been detected as in contact with the other protein in any of the snapshots were selected for the calculation. The snapshots shown in Fig. 1E are extracted by Gromacs module anaeig setting “nframes” as 5.

### Frustration calculation

Frustration analysis was conducted by Frustratometer 2^27^. All the residue in the BRD4:pairs refer to combination of highly, neutral and minimally frustration pairs. The PPI frustration level was defined by the number of frustration pairs divided by the number of all frustration pairs. In each frame, residue pairs on the PPI that any heavy atoms are in 4.5 Å were used to calculate the PPI frustration level. The NTD-BRD4 and CTD-BRD4 interface frustration number was analyzed separately. Protein structures were saved every 5ns from the simulation for the calculation.

### Residue contacts and residue-residue contact score

The protein-protein and protein ligand contacts were computed by python scripts GetContacts^41^. The contact polarity was determined from the atom-based contact types. Hydrogen bond, salt bridges and pi-cation interactions were considered polar. Pi-stacking van der Waals interactions are nonpolar. For each residue pair, if it contains polar atom interactions, it will be categorized as polar contacts. Otherwise, it is nonpolar contacts. Through the MD simulation, if over 10% of frames one residue pairs are polar, then will be colored as blue in the heatmap. Otherwise, it will be defined as nonpolar contact and colored black. Residue-residue contact score (RRCS) was utilized to evaluate the packing of constant contacts (>40%) on the protein-protein interface^28^. In each snapshot, the RRCS for one residue pair is the summation of distance-based score of all the heavy-atom pairs on both residues. To plot the RRCS heatmap, the score for each contact pair was averaged along all the frames.

### Water volume map, water number counting and water energy calculation

Water volmap analysis was conducted with VMD Volmap^42^. The calculation was conducted on all the frames that were overlayed to the first frame, and results were combined using the average of each snapshot. The map type was set to occupancy and resolution was set to 1 Å. The whole water molecule within 5 Å of protein complex were used for the calculation. The zoomed in map on PPI are shown in Figure 4. Water number near specific residues was counted by selecting water molecules within 3 Å of the heavy atoms. Water energy was calculated by WaterMap (version: v2022-2) panel in Maestro (Schrödinger, New York, NY, USA). To select the representative snapshots for water energy calculation, RMSD clustering on the four hydrophobic residues (F79, W81 on BRD4 and F102, P352 on cereblon) were carried out in Gromacs2022 with Gromos method and cutoff of 1 Å. The representative frame of top 4 cluster that covers at least 55% of the conformational population were saved. Water molecules that are within 7 Å of both cereblon and BRD4 at the same time were considered as PPI water and saved with the ternary complex for water energy calculation. The water energy state was defined by the excessed energy that was measured relative to bulk water. The water energy map of cluster 1 structure of L5 is shown in Figure 4.

### Kernel Density of the frustration level distribution calculations

The PPI frustration level distribution shown in Figure S8, was calculated using the average PPI frustration level from each MD trajectory (a total of 60 data points: 5 LDDs × 12 velocities). The bar graph showing the number of MD trajectories for any given value of the frustration level, was generated using the matplotlib “hist” function. For bar graph of all the data, the histogram bins were set to 20. For the bar graph of strong and weak LDDs separately, the bins for the weak and strong LDDs were set to 15 and 11, respectively, to ensure similar bar widths for both categories. The density curve was fitted using the “GaussianMixture” module from the “sklearn.mixture” package in Python, with three Gaussian components fitted to the distribution.

### Statistical analysis

The p-value was computed from T-test using scipy.stats.ttest_ind module in Python. If the p-value < 0.001, it would be labeled as “***”. If the p-value is between 0.001 and 0.01, it would be labeled as “**”. If the p-value is between 0.01 and 0.05, it would be labeled as “*”. If the p-value > 0.05, it would be labeled as “ns”.

## Acknowledgements

This work was funded by Bristol-Myers Squibb as sponsored research project to Vaidehi’s laboratory. We thank Dr. Supriyo Bhattacharya for the insightful discussions on protein frustration calculations.

## Supporting Information

**Figure S1.**
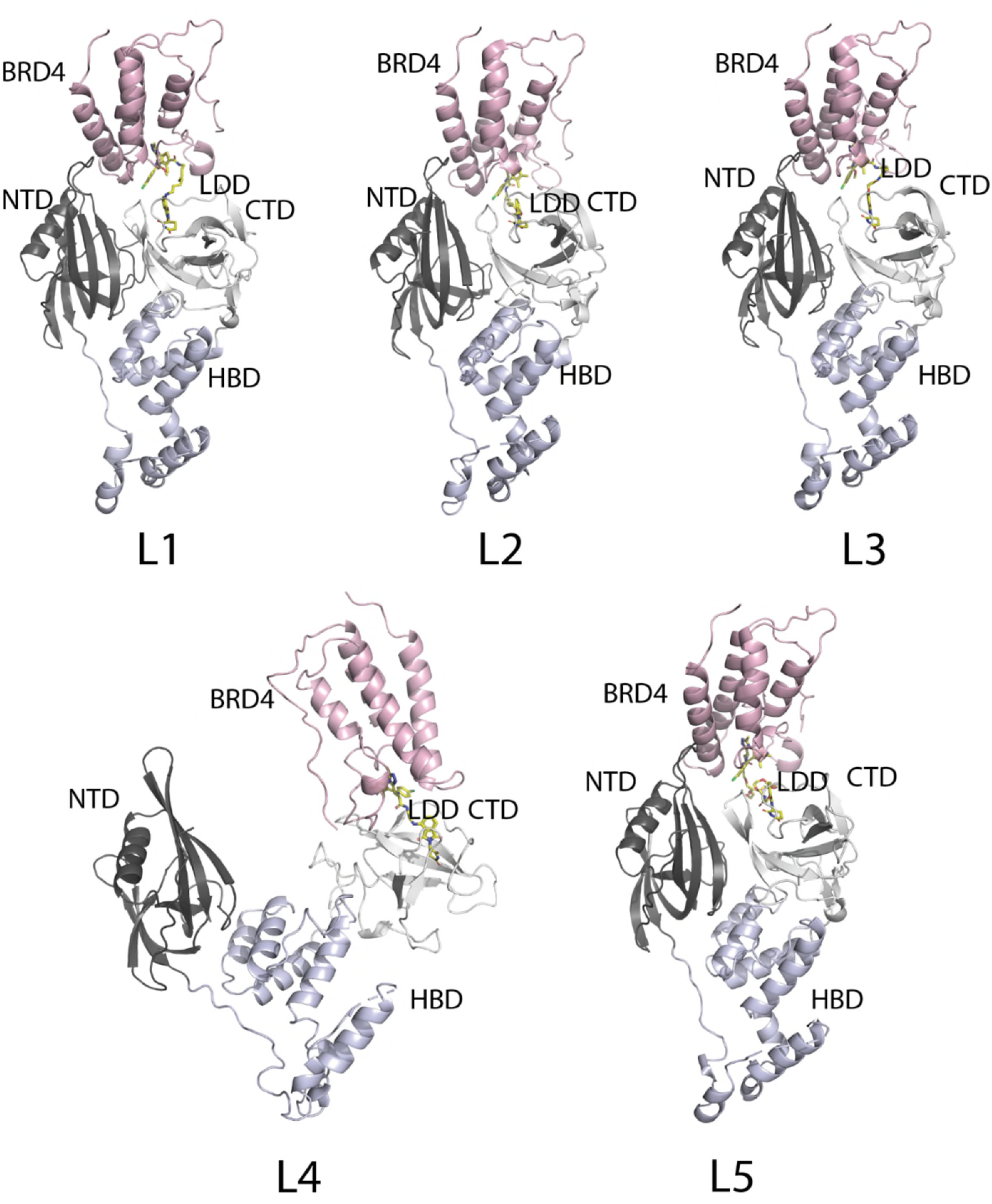
The ternary complex of L1(6BN9), L2(6BOY), L3(6BN7), L4(6BNB), L5(6BN8). The LDD molecule of L1 was modeled using L3 as template. The LDD molecule of L4 and L5 were modeled using L2 as template. The BRD4 in L4 complex adopts a different orientation than L1, L2, L3, L5.

**Figure S2.**
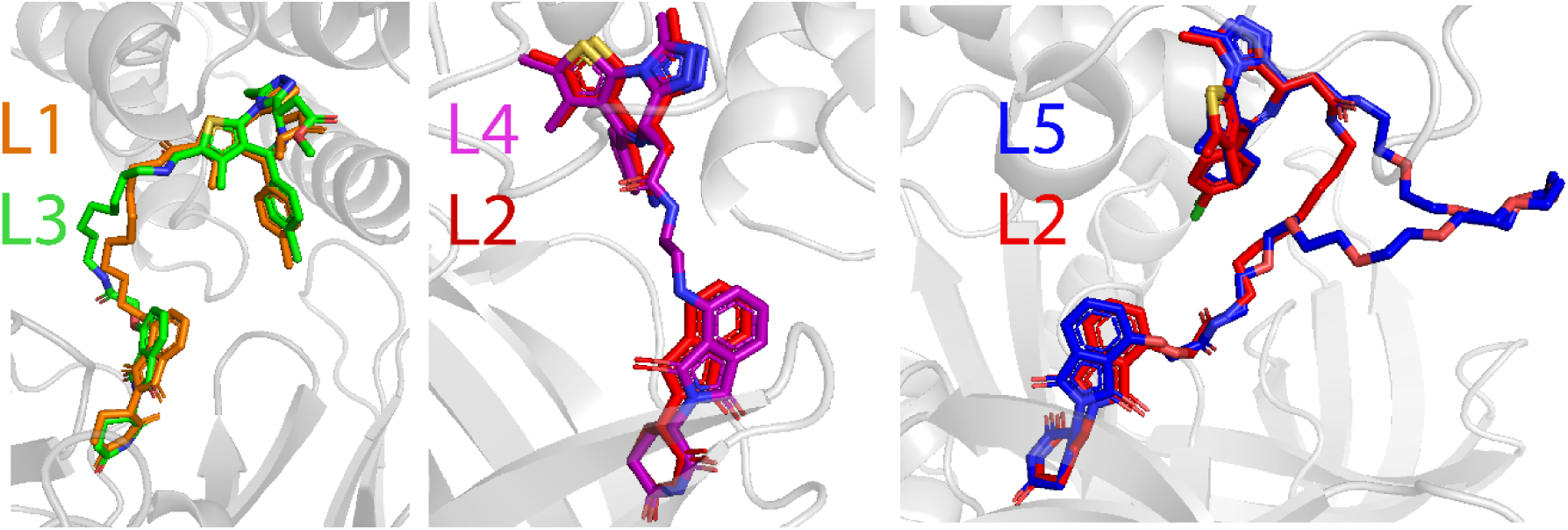
Structure figure of homology modeling of the linker. B) RMSD in coordinates of the residues in the warhead binding site for all the five LDDs.

**Figure S3.**
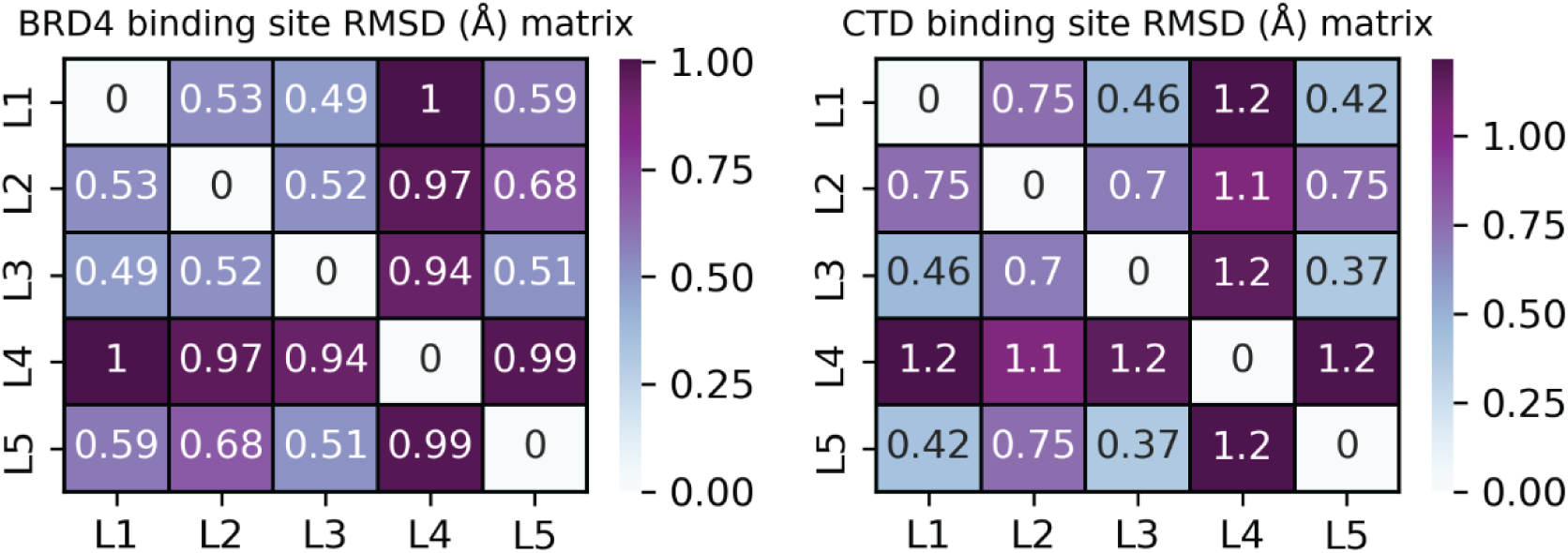
RMSD in coordinates of the residues in the warhead binding site for all the five LDDs.

**Figure S4.**
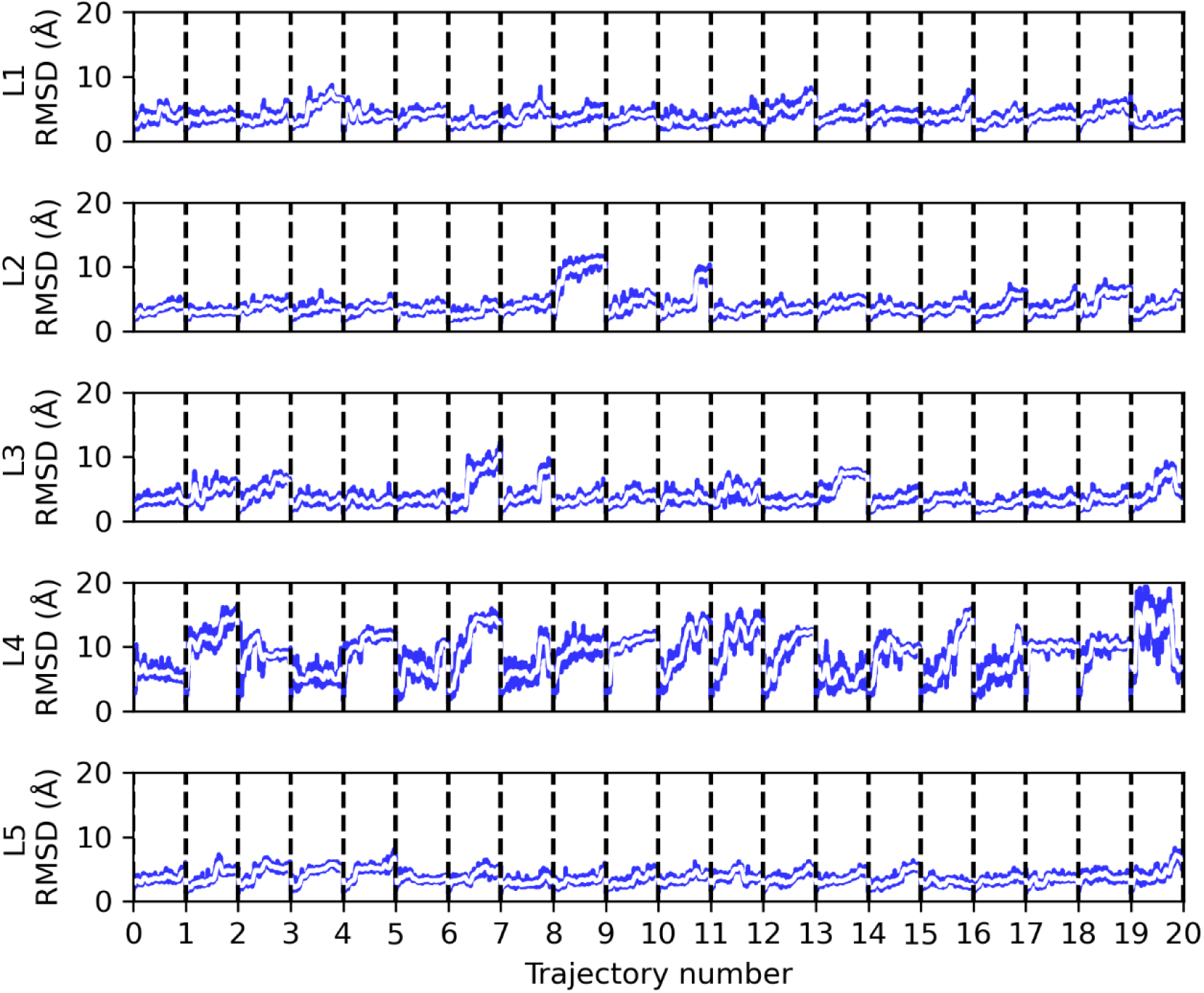
RMSD over time of the five LDD ternary complexes from the MD simulations.

**Figure S5.**
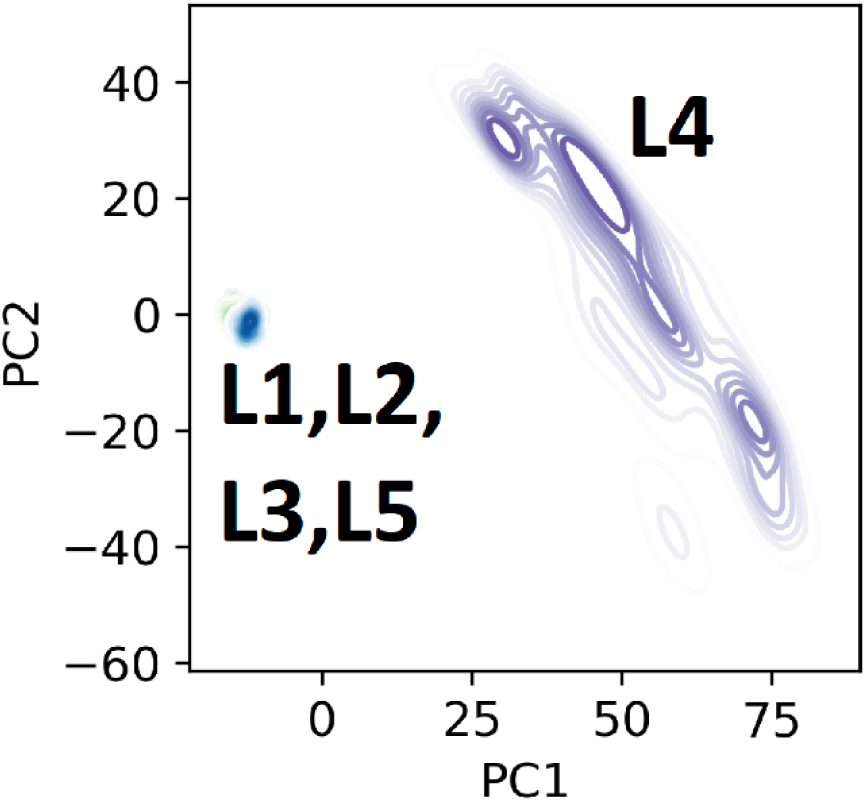
PC landscape of PC1 and PC2 when combine all the five LDD complex together, L4 motion dominant the PC space.

**Figure S6.**
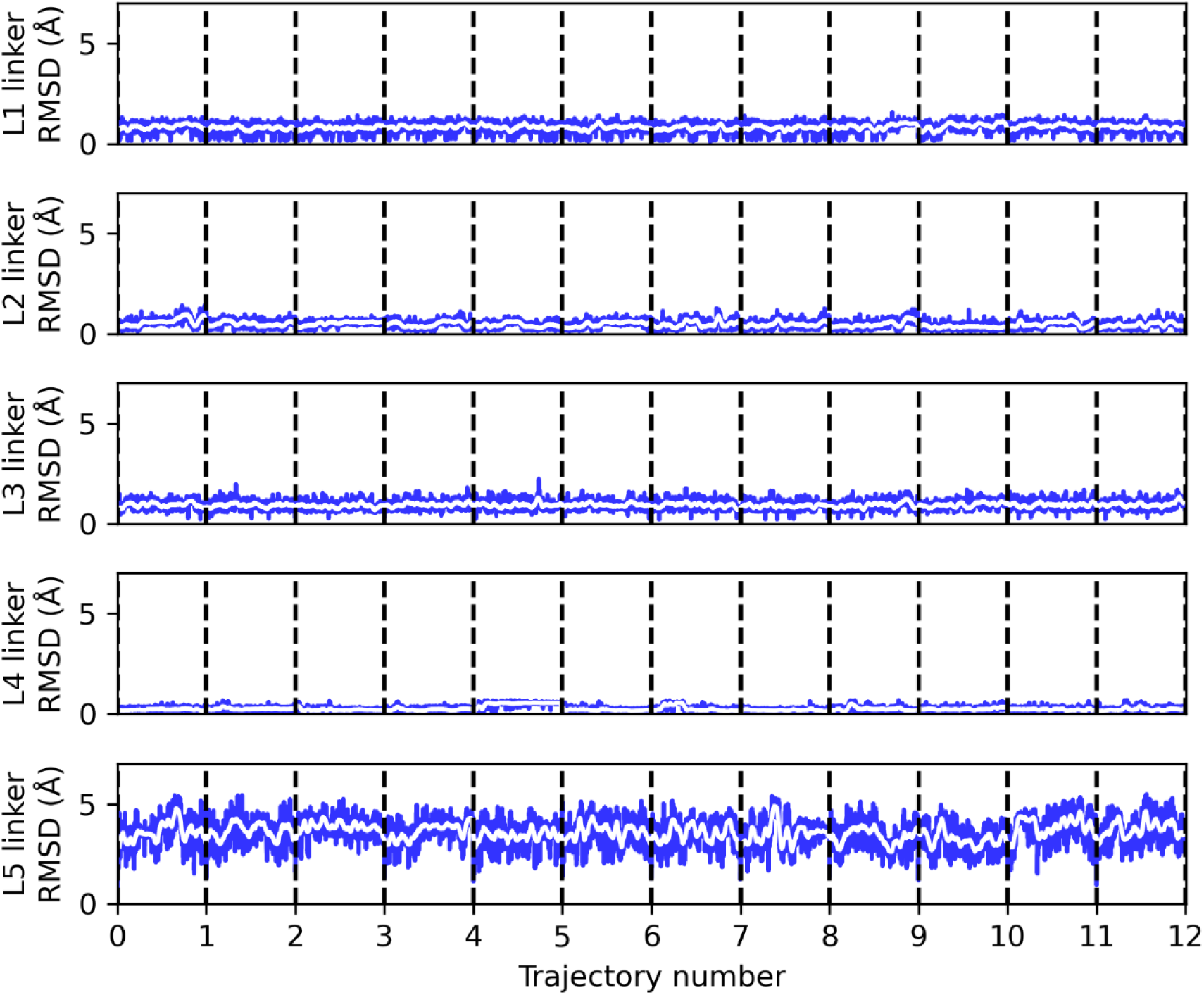
LDD linker RMSD over time from the five LDD complex MD simulations.

**Figure S7.**
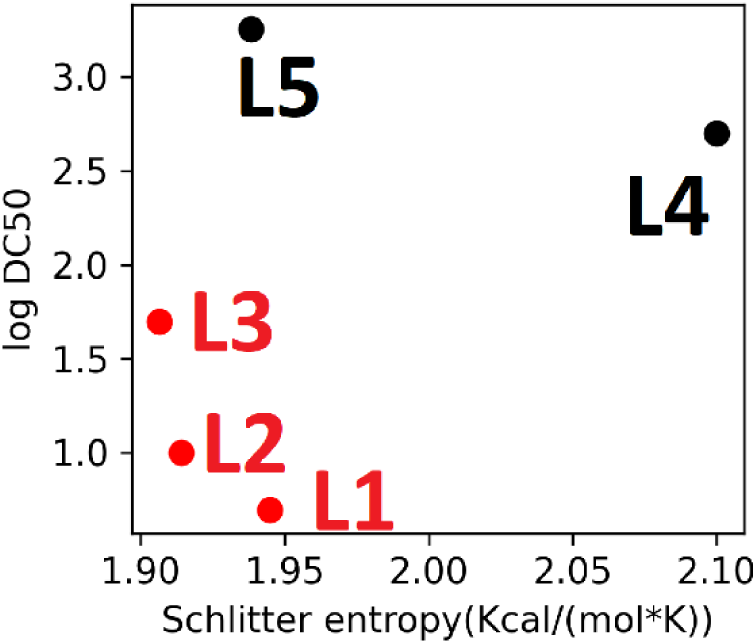
Global conformational entropy of the five LDD complexes.

**Figure S8.**
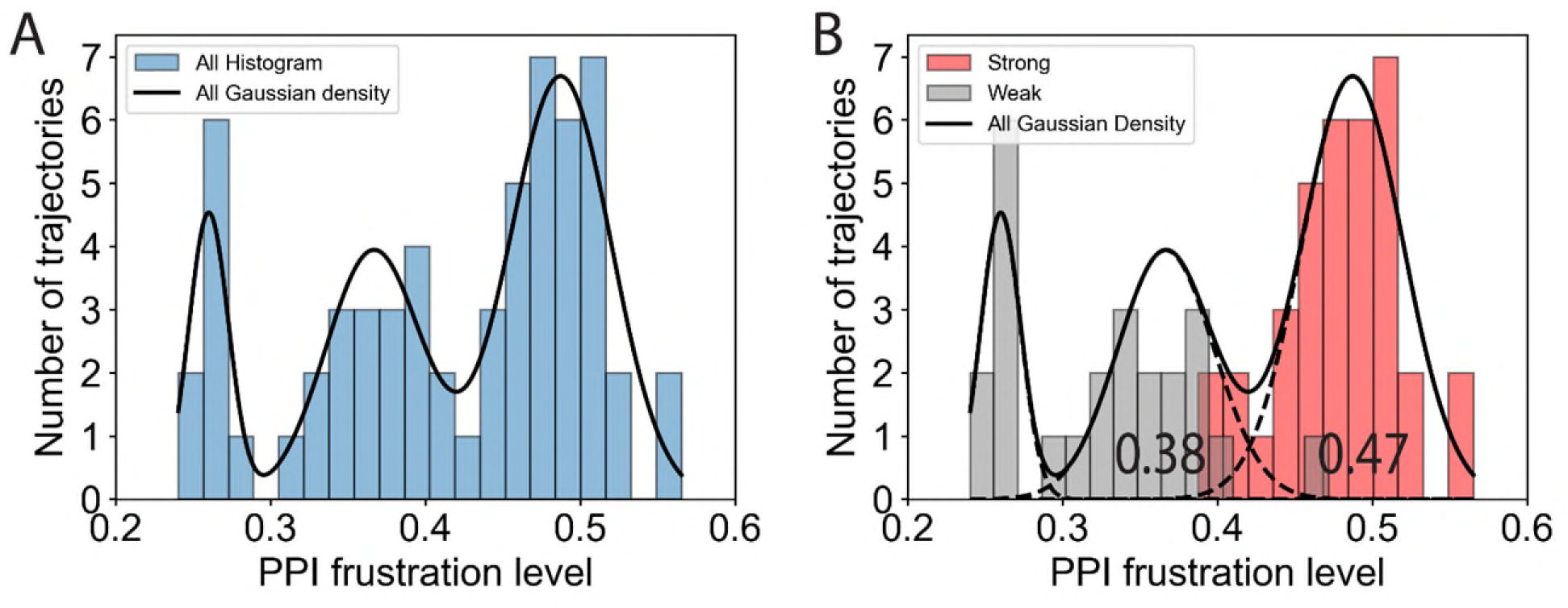
The distribution of average PPI frustration level in each trajectory. A) The distribution includes all trajectories, encompassing both strong and weak LDDs. Three Gaussian kernels are fitted to the overall distribution. B) Trajectories for weak LDDs (gray) and strong LDDs (red) are plotted separately. The Gaussian kernels are extended as dashed lines to intersect with the x-axis. The right intersection of the second Gaussian is at 0.38, while the left intersection of the third Gaussian is at 0.47.

**Figure S9.**
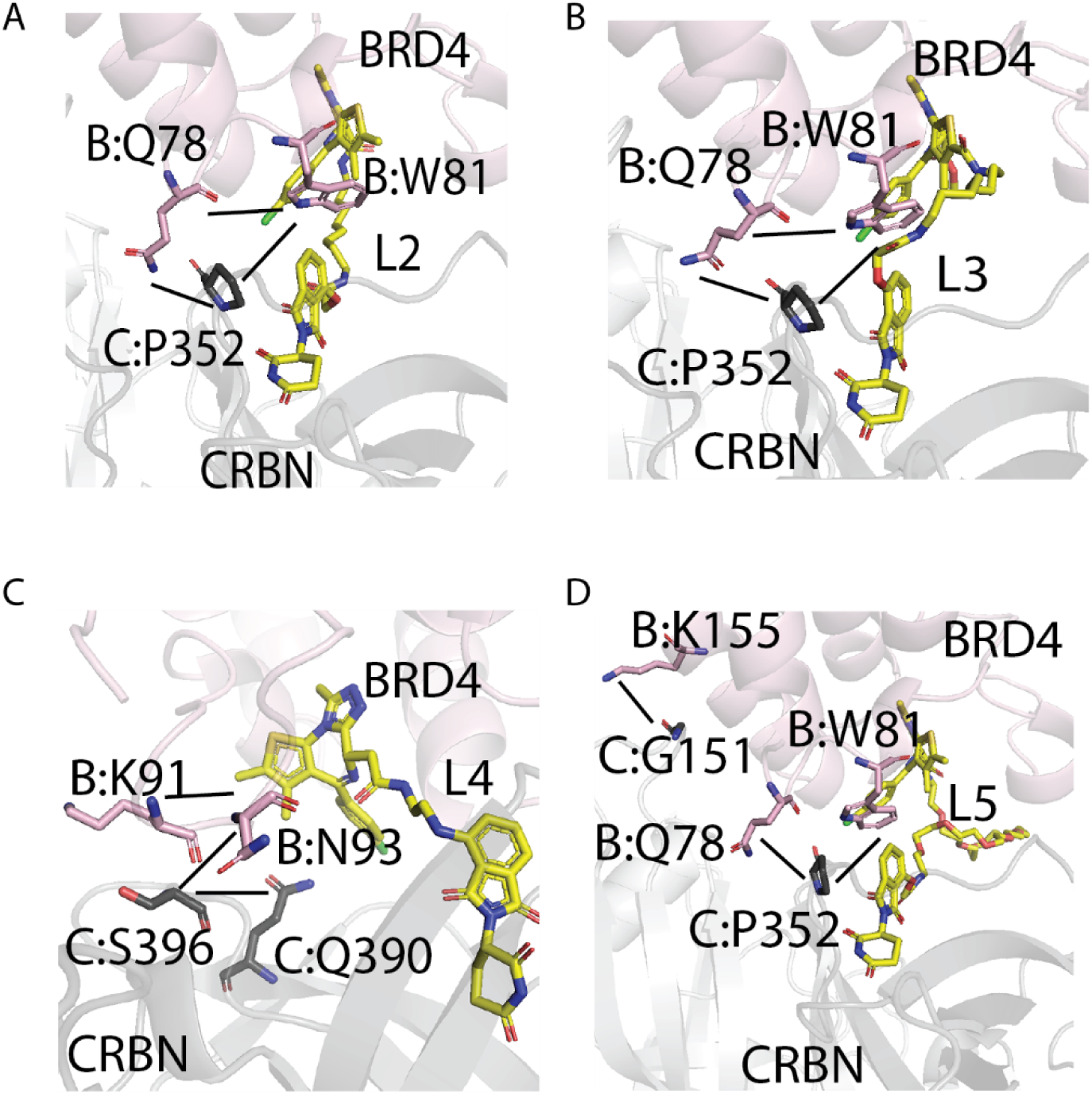
The top 3 persistent (in terms of frequency in MD simulations) highly frustrated contact pairs in L2-L5 (A-D) mapped on the 3D structure. The short black lines show the persistent highly frustrated contact pairs. BRD4’s cartoon and sidechains are in pink. Cereblon’s cartoon and sidechains are in grey.

**Figure S10.**
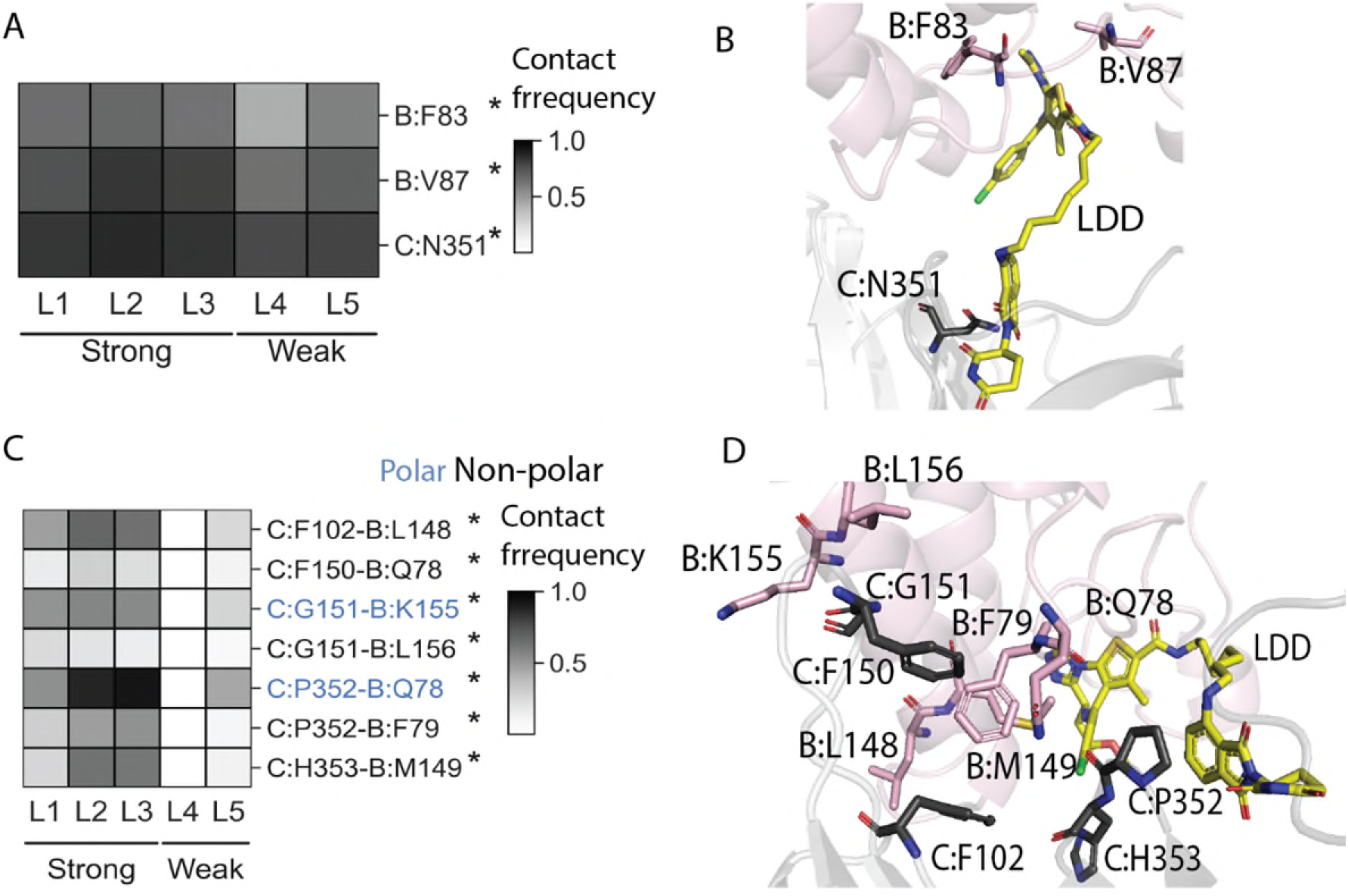
A) Heatmap showing the contact frequency between LDD’s binding moiety and protein residues. Here protein includes BRD4(B) and cereblon(C). B) 3D structure highlighting the residues in the LDD binding moiety-protein residues contact heatmap. C) Heatmap showing the contact frequency between BRD4(B) and cereblon(C). D) 3D structure highlighting the residues shown in the PPI contact heatmap.

**Figure S11.**
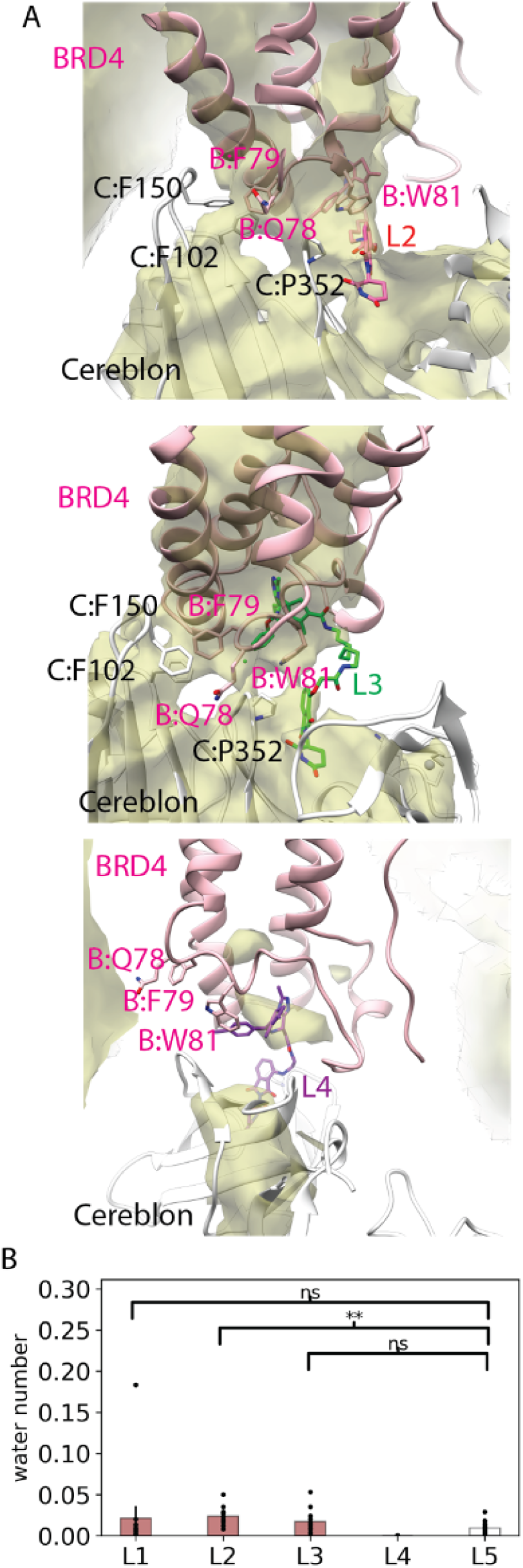
Hydrophobic region in BRD4-Cereblon PPI. A) Water occupancy map for L2, L3 and L4 complex. The water occupancy < 1% region is shown as yellow cloud. B) Number of water in the C:F102-B:F79-C:F150 hydrophobic patch. P-value for significance: L1-L5: 0.22; L2-L5: 0.001; L3-L5: 0.06.

