## supplemental figures for "Insights From Protein Frustration Analysis of BRD4-Cereblon Degrader Ternary Complexes Show Separation of Strong from Weak Degraders"

[3] Bristol-Myers-Squibb

[4] Present Address: Septerna, 250 E Grand Ave, South San Francisco, CA 94080

**Supporting Information**

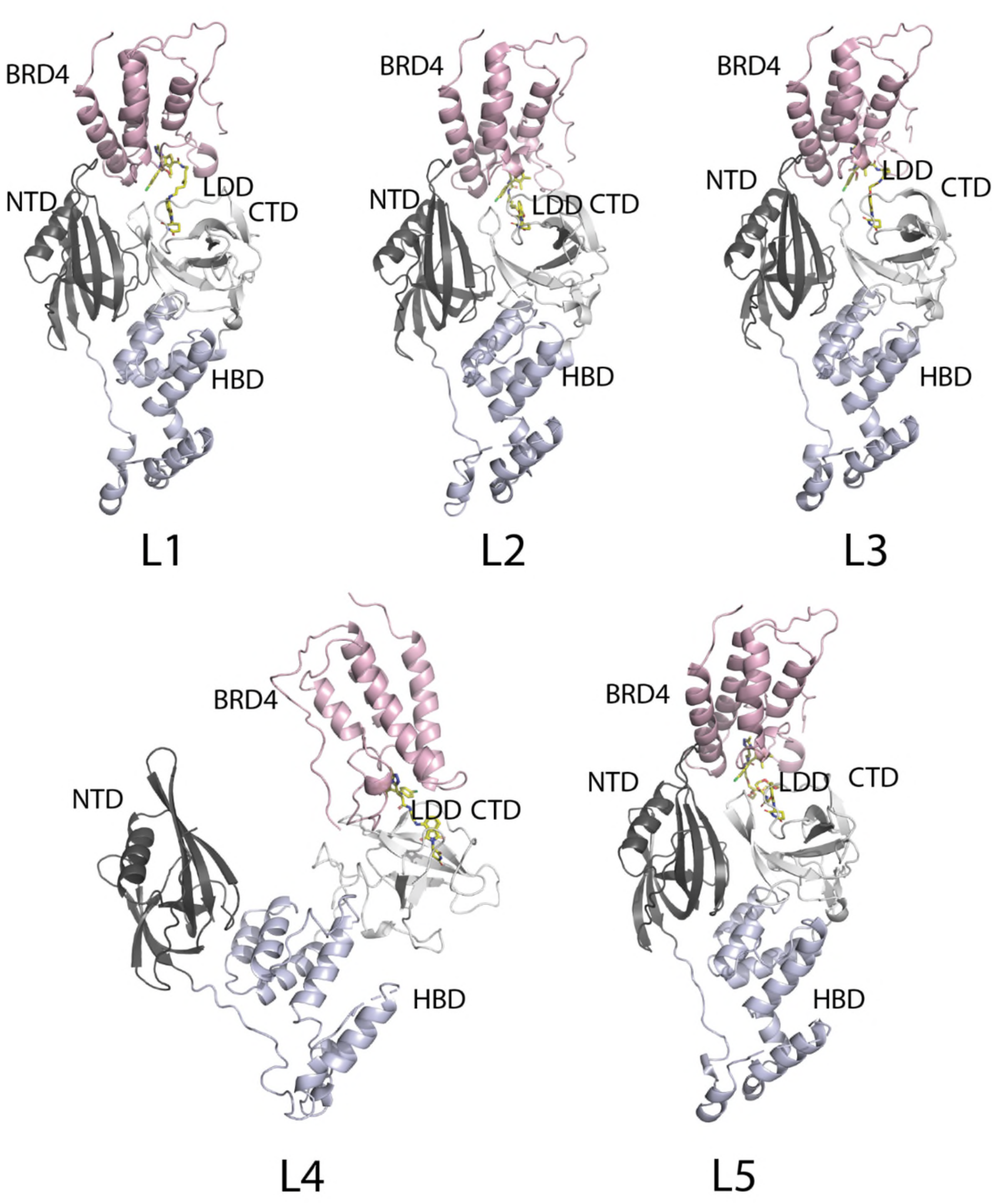


**Figure S1.** The ternary complex of L1(6BN9), L2(6BOY), L3(6BN7), L4(6BNB), L5(6BN8). The LDD molecule of L1 was modeled using L3 as template. The LDD molecule of L4 and L5 were modeled using L2 as template. The BRD4 in L4 complex adopts a different orientation than L1, L2, L3, L5.

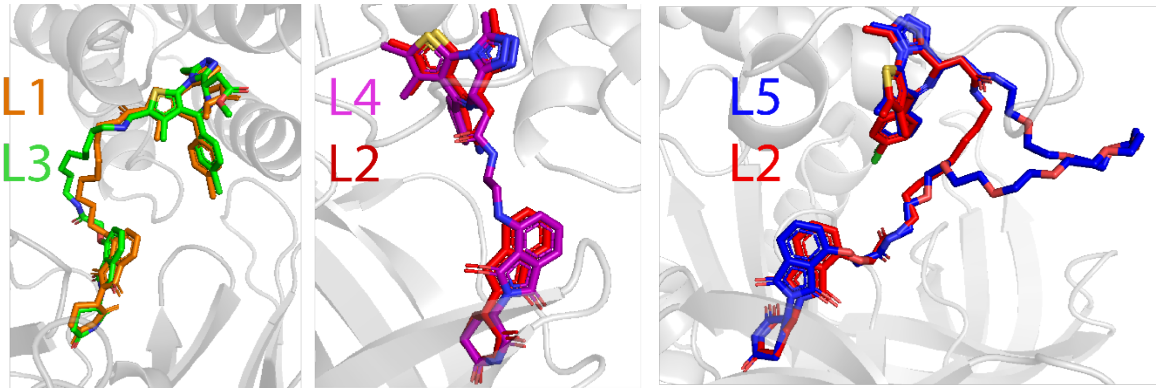


**Figure S2.** Structure figure of homology modeling of the linker. B) RMSD in coordinates of the residues in the warhead binding site for all the five LDDs.

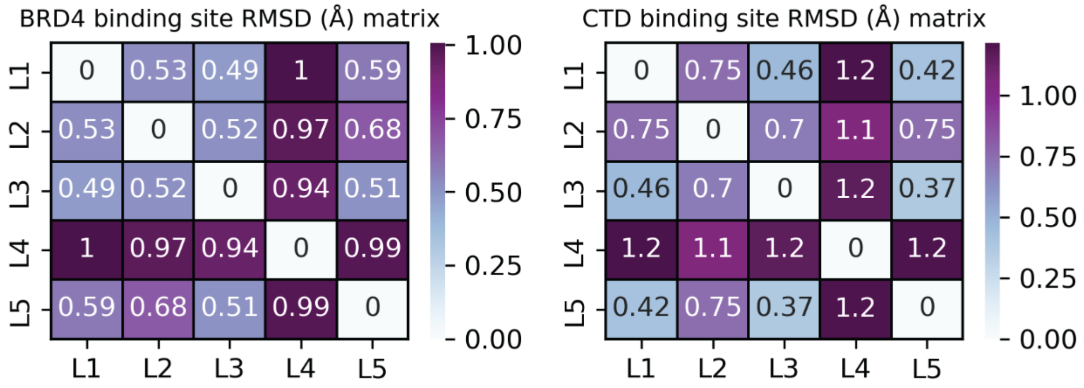


**Figure S3.** RMSD in coordinates of the residues in the warhead binding site for all the five LDDs.

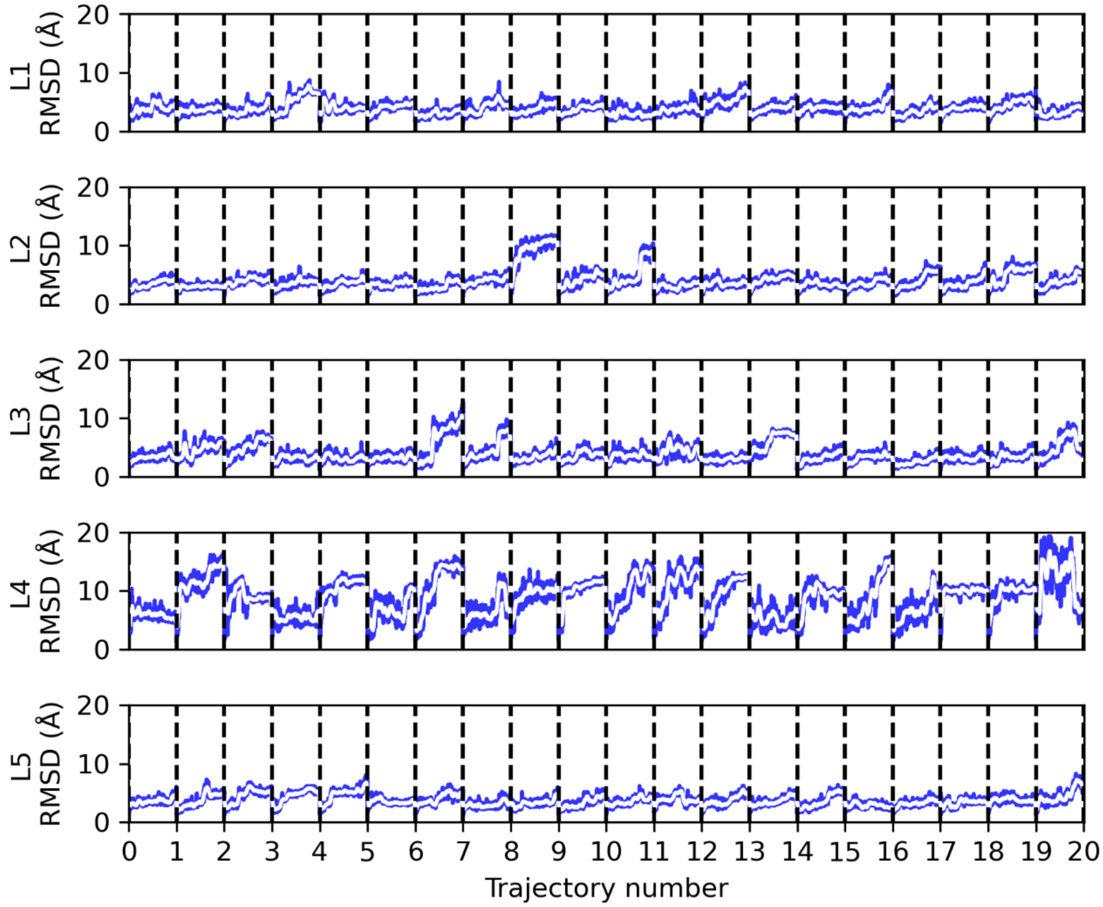


**Figure S4.** RMSD over time of the five LDD ternary complexes from the MD simulations.

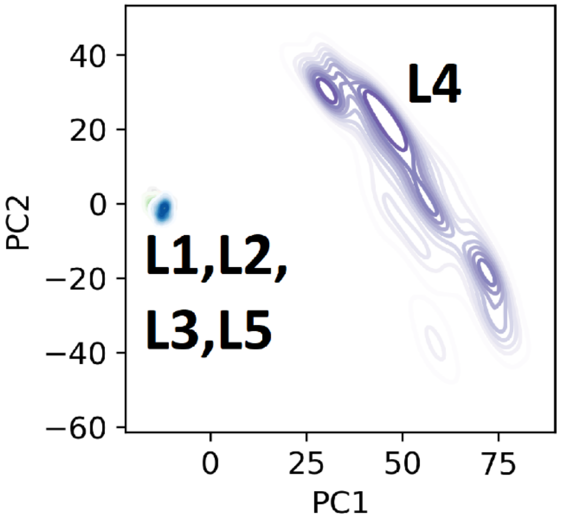


**Figure S5.** PC landscape of PC1 and PC2 when combine all the five LDD complex together, L4 motion dominant the PC space.

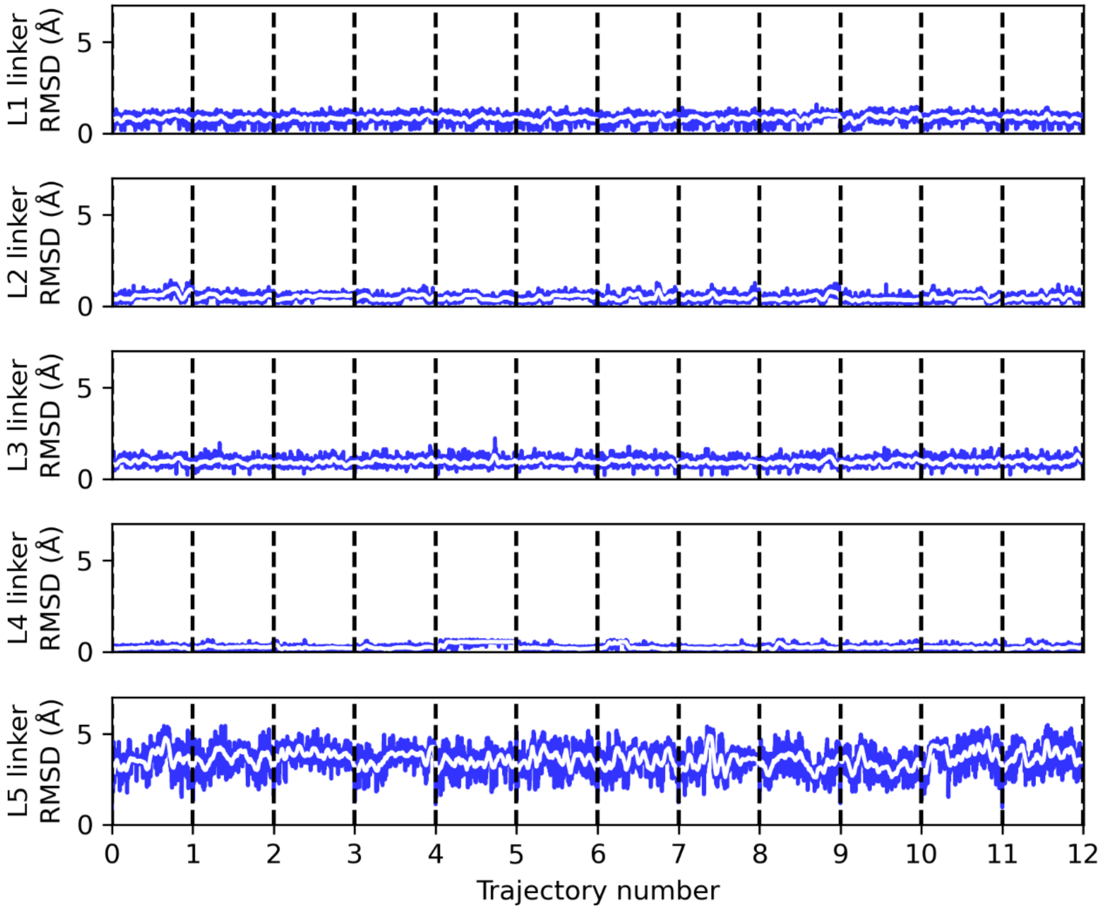


**Figure S6.** LDD linker RMSD over time from the five LDD complex MD simulations.

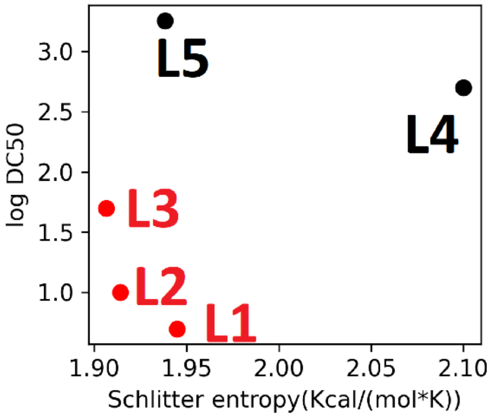


**Figure S7.** Global conformational entropy of the five LDD complexes.

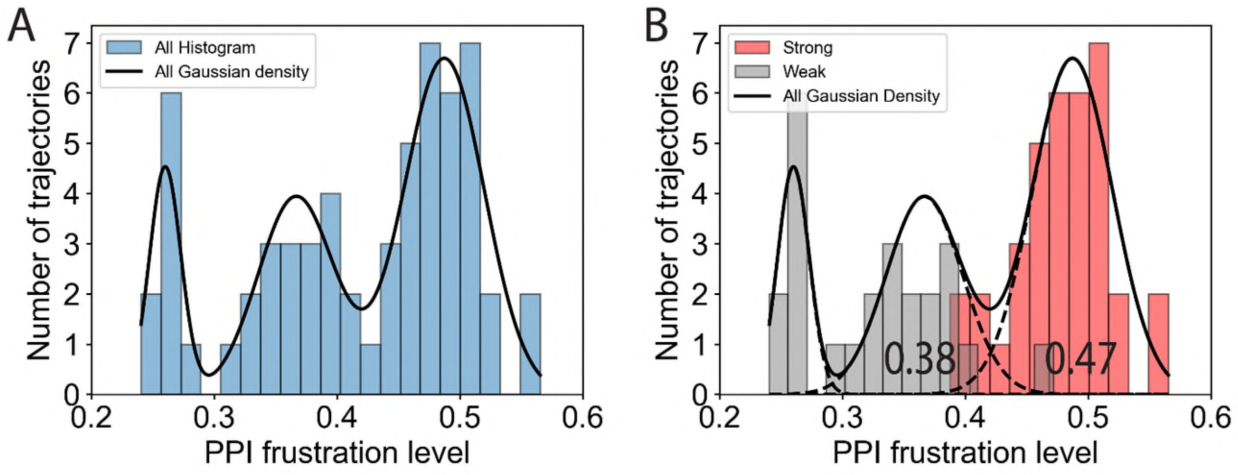


**Figure S8.** The distribution of average PPI frustration level in each trajectory. A) The distribution includes all trajectories, encompassing both strong and weak LDDs. Three Gaussian kernels are fitted to the overall distribution. B) Trajectories for weak LDDs (gray) and strong LDDs (red) are plotted separately. The Gaussian kernels are extended as dashed lines to intersect with the x-axis. The right intersection of the second Gaussian is at 0.38, while the left intersection of the third Gaussian is at 0.47.

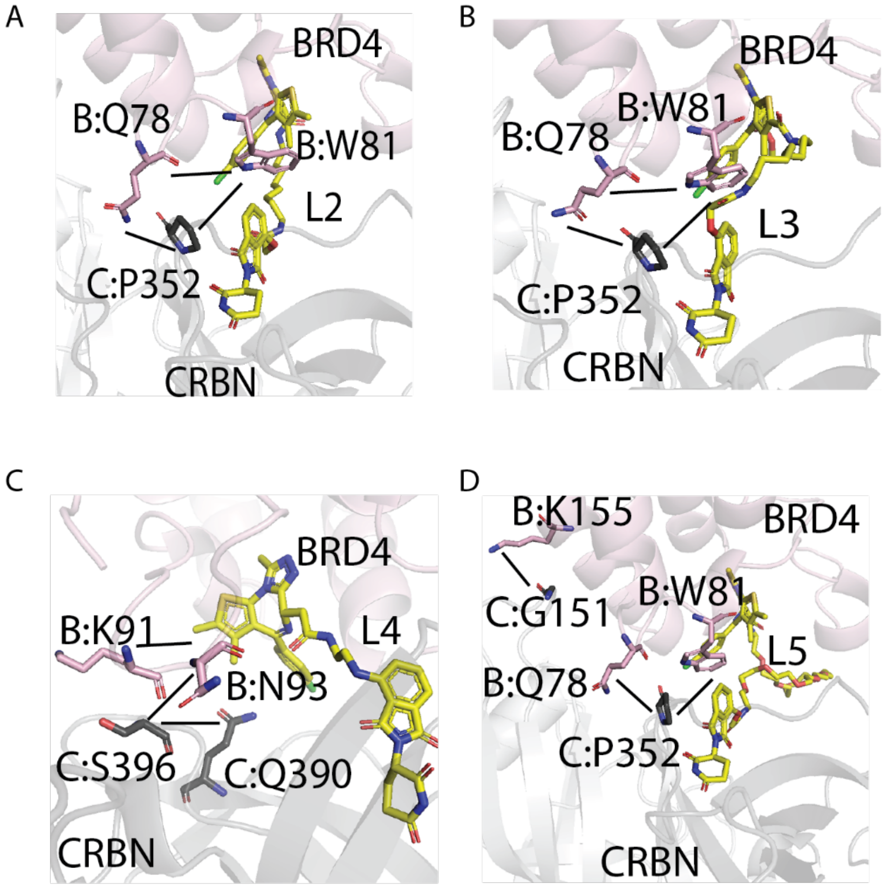


**Figure S9.** The top 3 persistent (in terms of frequency in MD simulations) highly frustrated contact pairs in L2-L5 (A-D) mapped on the 3D structure. The short black lines show the persistent highly frustrated contact pairs. BRD4's cartoon and sidechains are in pink. Cereblon's cartoon and sidechains are in grey.

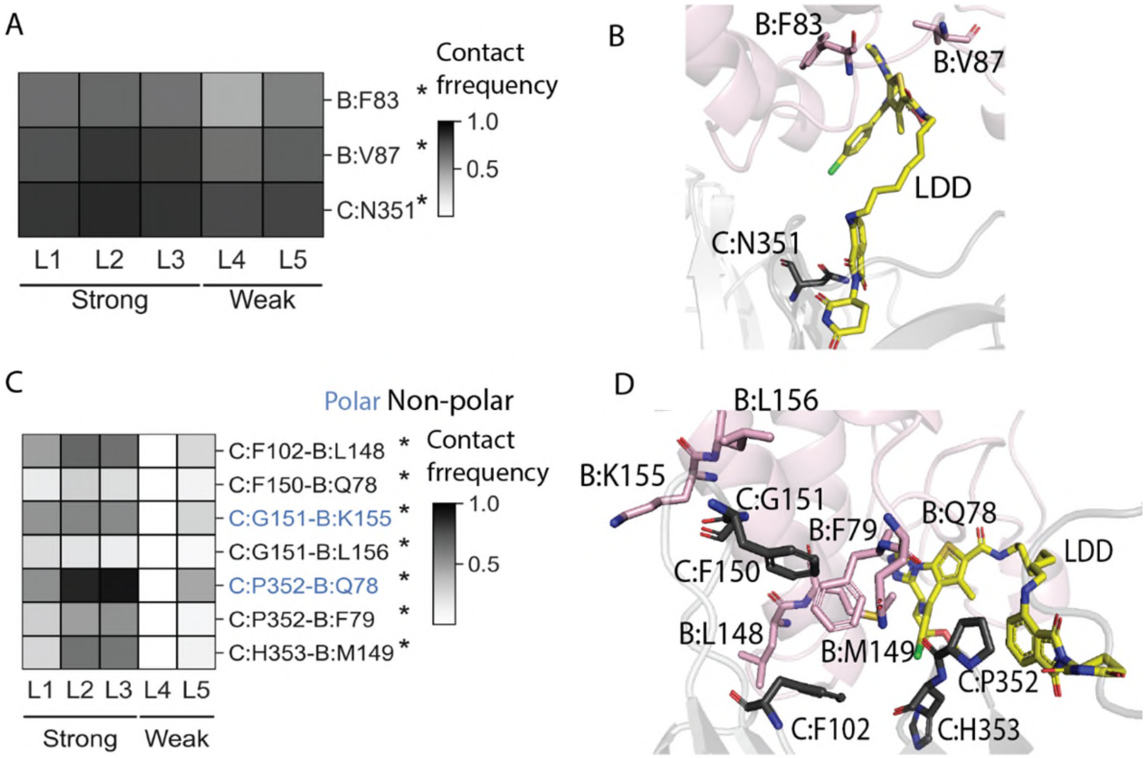


**Figure S10.** A) Heatmap showing the contact frequency between LDD's binding moiety and protein residues. Here protein includes BRD4(B) and cereblon(C). B) 3D structure highlighting the residues in the LDD binding moiety-protein residues contact heatmap. C) Heatmap showing the contact frequency between BRD4(B) and cereblon(C). D) 3D structure highlighting the residues shown in the PPI contact heatmap.

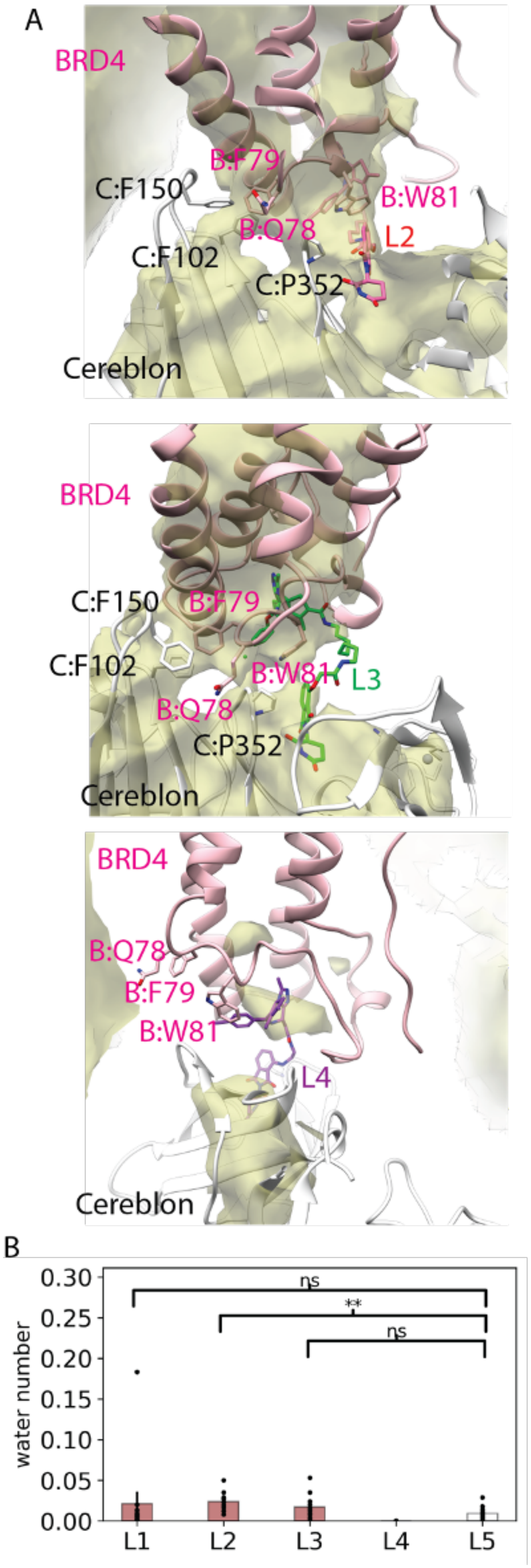


**Figure S11.** Hydrophobic region in BRD4-Cereblon PPI. A) Water occupancy map for L2, L3 and L4 complex. The water occupancy < 1% region is shown as yellow cloud. B) Number of water in the C:F102-B:F79-C:F150 hydrophobic patch. P-value for significance: L1-L5: 0.22; L2-L5: 0.001; L3-L5: 0.06.
